# Identification of amyloid beta oligomers in locus coeruleus (LC) neurons of Alzheimer’s patients and their impact on LC oxidative stress, inhibitory neurotransmitter receptors and neuronal excitability

**DOI:** 10.1101/2020.05.14.096396

**Authors:** Louise Kelly, Mohsen Seifi, Ruolin Ma, Scott Mitchell, Uwe Rudolph, Kirsten L. Viola, William L. Klein, Jeremy J. Lambert, Jerome D. Swinny

**Affiliations:** School of Pharmacy & Biomedical Sciences, University of Portsmouth, Portsmouth, PO12DT, United Kingdom; Neuroscience, Division of Systems Medicine, Ninewells Hospital & Medical School, Dundee University, Dundee, DD19SY, Scotland, United Kingdom; Department of Comparative Biosciences, College of Veterinary Medicine, and Carl R. Woese Institute for Genomic Biology, University of Illinois at Urbana-Champaign, Urbana, IL, USA; Department of Neurobiology and Physiology, Northwestern University, Evanston, Illinois, USA

**Keywords:** dementia, noradrenaline, GABA-A receptors, glycine transporters, stress

## Abstract

Amyloid β oligomers (AβO) are potent modulators of two key Alzheimer’s pathological processes, namely synaptic dysfunction and tau tangle formation in various brain regions. Remarkably, the impact of AβO in one of the earliest brain regions to exhibit Alzheimer’s pathology, the locus coeruleus (LC), remains to be determined. Of particular importance is the effect of AβO on the excitability of individual LC neurons. This parameter determines brain-wide noradrenaline (NA) release, and thus NA-mediated brain functions, including cognition, emotion and immune function, which are all severely compromised in Alzheimer’s. Using a mouse model of increased Aβ production (APP-PSEN1), together with correlative histopathological analyses in post mortem Alzheimer’s patient samples, we determined the impact of Aβ pathology on various correlates of LC neuronal integrity. AβO immunoreactivity in the LC of APP-PSEN1 mice was replicated in patient samples, presenting as individual clusters located both intraneuronally, in mitochondrial compartments, as well as extracellularly in association with inhibitory synapses. No specific signal was detected in either patient control or wild type mouse samples. Accompanying this AβO expression profile was LC neuronal hyperexcitability and indicators of oxidative stress in APP-PSEN1 mice. LC hyperexcitability arose from a diminished inhibitory effect of GABA, due to impaired expression and function of the GABA-A receptor (GABA_A_R) α3 subunit. Importantly, this altered LC α3-GABA_A_R expression profile overlapped with AβO expression in both APP-PSEN1 mice and Alzheimer’s patient samples. Finally, strychnine-sensitive glycine receptors (GlyRs) remained resilient to AβO-induced changes and their activation reversed LC hyperexcitability. Alongside this first demonstration of AβO expression in the LC of Alzheimer’s patients, the study is also first to reveal a direct association between AβO and LC neuronal excitability. GlyR-α3-GABA_A_R modulation of AβO-dependent LC hyperexcitability could delay the onset of cognitive and psychiatric symptoms arising from LC-NA deficits, thereby significantly diminishing the disease burden for Alzheimer’s patients.

## Introduction

Synaptic dysfunction is a key pathological mechanism underlying the mild cognitive, memory and neuropsychiatric impairments that typify the earliest stages of Alzheimer’s (de Wilde et al. 2016; DeKosky and Scheff 1990; Spires-Jones and Hyman 2014). Given the brain’s established propensity for synaptic remodelling, identifying the molecular and cellular mechanisms of such synaptic pathology, prior to the onset of irreversible neuronal death, likely represents one of the best interventionist strategies against this disease (Colom-Cadena et al. 2020; Jackson et al. 2019). Convergent lines of evidence point to a crucial role for the soluble, oligomeric variants of amyloid β (AβO) (Lambert et al. 1998; Viola and Klein 2015) in mediating such synaptic dysregulation (Ding et al. 2019; Koffie et al. 2009).

Experimentally, AβO bind directly to synaptic junctions (Koffie et al. 2012; Koffie et al. 2009; Lacor et al. 2004), alters excitatory and inhibitory synaptic transmission (Calvo-Flores Guzman et al. 2020; He et al. 2019; Marcantoni et al. 2020), the expression and location of synaptic neurotransmitter receptors (Gu et al. 2009; Ulrich 2015) as well influencing synaptic plasticity (Calvo-Flores Guzman et al. 2020) and learning and memory (Lee et al. 2006). This association is also evident in humans, with a strong association demonstrated between AβO and dementia (Bao et al. 2012; Mc Donald et al. 2010). While such compelling evidence exist for AβO pathology in cortical brain regions, less is known about the impact of this pathway in one of the first regions to exhibit quintessential Alzheimer’s histopathology, namely the locus coeruleus (LC) (Braak et al. 2011; Weinshenker 2018). In particular, we do not know whether AβO-related pathology alters the core determinant of LC-mediated brain functions, the spontaneous neuronal firing rate, and if so, through which mechanisms?

The relevance of LC neurobiology to the understanding of the spectrum Alzheimer’s-related symptoms is underscored by the overlap between the brain functions this system modulates, and those which are impaired in this condition. These include cognition (Sara 2015), arousal (Carter et al. 2010), emotion (Itoi and Sugimoto 2010) and the stress response (Valentino and Van Bockstaele 2008). The coordinated brain-wide release of the LC’s signature neurotransmitter noradrenaline (NA), and thus LC-NA mediated brain functions, arises from the complex firing patterns of LC neurons, which are brain (Usher et al. 1999), behaviour (Carter et al. 2010; McCall et al. 2015) and disease-state (Sanchez-Padilla et al. 2014) dependent. This emphasises the importance of understanding the molecular machinery of the LC and how this contributes to regulating LC neuronal excitability in health and disease.

The focus of Alzheimer’s pathology within the LC has been predominantly on hyperphosphorylated tau tangles (Braak et al. 2011) because of their early presence during the disease spectrum. In contrast, Aβ-related pathology has largely been overlooked because only insoluble Aβ-containing plaques have been documented within the LC, and they present only in the final stages of the condition (Thal et al. 2002). However, neither the presence of the more toxic soluble AβO within LC, nor their effects on LC neuronal excitability, has been demonstrated. This is important because of the evidence indicating that AβO, rather than Aβ plaques, mediate the neuro-toxic effects of the Aβ pathway (Cline et al. 2018; Ghag et al. 2018). Of particular relevance to the LC, and its early presentation of Alzheimer’s pathology, is the emerging synergistic role of AβO in tau tangle formation (Pickett et al. 2019; Shin et al. 2019b). Intriguingly, while notable LC neuronal loss, attributed to tauopathy, is a core feature in Alzheimer’s (Chan-Palay and Asan 1989), there appears to be discordance between the onset of such pathology, and LC neurodegeneration (Weinshenker 2018). Given such considerable evidence, it is reasonable to speculate that in the LC, initial, pre-symptomatic insults in the form of tau tangle and AβO interactions, cooperate to set in motion a range of pathological changes such as synaptic dysfunction, oxidative stress (De Felice et al. 2007) and altered LC excitability (Sanchez-Padilla et al. 2014). Such changes are likely to prove crucial in terms of the long-term viability of cells not only within the LC, but also other brain regions because impaired LC integrity directly influences the progression of Alzheimer’s pathology in cortical regions (Heneka et al. 2010; Heneka et al. 2006). To begin to explore the potential influence of AβO on the LC-NA system, and thus connected brain regions, in the current study, we investigate the impact of Aβ, and AβO in particular, on the molecular and physiological correlates of the LC.

## Materials and Methods

### Ethics

All procedures involving animal experiments were approved by the Animal Welfare and Ethical Review Body of the University of Portsmouth and were performed by a personal license holder, under a Home Office-issued project license, in accordance with the Animals (Scientific Procedures) Act, 1986 (UK) and associated procedures.

### Animals

The B6C3-Tg (APPswe, PSEN1dE9) double transgenic mouse model, that results in accelerated production of Aβ (Borchelt et al. 1997), termed APP-PSEN1 onwards, was the main strain used in the study. APP-PSEN1 mice exhibit a loss of LC neurons (O’Neil et al. 2007) and their efferent projections (Liu et al. 2008), in old age (> 12 months). However, the expression of AβO, together with the changes in native neurotransmitter-receptors and physiology of individual LC neurons have yet to be determined, especially in young mice. Therefore, animals between 2 and 4 months of age were used unless otherwise specified. Breeding pairs were obtained from The Jackson Laboratory (USA). A colony was established at the University of Portsmouth by crossing transgenic mice with C57BL/6J mice. In all experiments, wild type (WT) littermates were used as controls for APP-PSEN1 subjects. In a subset of experiments, GABA_A_R α3 subunit gene deleted (α3^−/−^) mice and their WT littermates (Yee et al. 2005), bred on a C57BL/6J background, as well as mice expressing green fluorescent protein (GFP) under the promotors of the astroglial cytoskeletal protein, glial fibrillary acidic protein (GFP-GFAP) (Lalo et al. 2006) were used. Male mice were used throughout all experiments.

Mice were group housed, given free access to water and standard rodent chow (Special Diet Service U.K.), maintained on a 12 h alternating light-dark regimen with lights on at 7–7:30 a.m. The temperature and the humidity were controlled at 21 ± 2 °C and 50 ± 5% respectively. Experiments were conducted on mice obtained from the first two generations of WT and mutant breeding pairs.

Data were reproduced from at least three different litters. Measures to optimise reproducibility and minimise bias included randomisation, in terms of genotype and litters, when selecting animals for experiments, as well as blinding the investigators at various stages.

### Human samples

Formaldehyde-fixed, paraffin embedded, 8 µm thickness brain sections containing the LC were provided by Brains for Dementia Research tissue banks, from n = 5 control and 5 Alzheimer’s subjects. Demographic details of the subjects are detailed in SI Table 2.

### Immunohistochemistry: Animal Tissue

It should be noted that since the main priority was to investigate the association of Aβ expression with synaptic, and therefore integral membrane proteins, animal tissue preparation was optimised to reveal such epitopes by using a comparatively weak fixation protocol (Lorincz and Nusser 2008a; Lorincz and Nusser 2008b).Experiments were conducted according to previously described protocols (Corteen et al. 2011). Briefly, mice were anaesthetised with isoflurane and then pentobarbitone (1.25 mg/kg of bodyweight (i.p), perfused transcardially with 0.9% saline solution for 2 minutes followed by 12 minutes with a fixative solution consisting of 1% freshly depolymerised formaldehyde and 15% v/v saturated picric acid in 0.1 M phosphate buffer (PB), pH 7.4. Brains were dissected and post-fixed over night at room temperature in the same perfusion fixative, after which 60 μm-thick horizontal sections of the LC were prepared using a vibratome (Leica VT 1000). The sections were washed in 0.1 M PB to remove any residual fixative and stored in 0.1 M PB containing 0.05% sodium azide. All washing steps were performed three times for 10 minutes using TRIS-buffered saline containing 0.3% Triton X100 (TBS-Tx). For the immunolocalisation of GlyRs, a proteolytic antigen retrieval method was used according to previous protocols (Corteen et al. 2011). Non-specific binding of secondary antibodies was minimised by incubating the tissue sections in a TBS-Tx solution containing 20% normal serum from the species of which the secondary antibodies were raised (Vector Laboratories, Burlingame, CA, USA), for two hours at room temperature, followed by incubation in a cocktail of primary antibodies overnight at 4°C (see SI Table 1 for details of primary antibodies used). The next day, the sections were washed in TBS-Tx, then incubated at room temperature in a cocktail of appropriate secondary antibodies, conjugated with Alexa Fluor 488, indocarbocyanine (Cy3) or indocarbocyanine (Cy5), (Jackson ImmunoResearch), for two hours before being washed, mounted on glass slides, air dried and coverslipped using Vectashield mounting medium (H-100, Vector Laboratories). In a subset of experiments using animals older than 6 months, sections were washed and incubated with Autofluorescence Eliminator (Millipore) according to the manufacturer’s protocol, before being cover-slipped.

### Immunohistochemistry: Human Tissue

Paraffin embedded brain sections containing the LC of 8 µm thickness were provided by Brains for Dementia Research tissue banks, from n = 5 control and 5 AD subjects. Demographic details of the subjects are detailed in SI Table 2. All washing steps were performed three times for 5 min using 0.1M PB. Sections were first deparaffinised and then rehydrated through a graded series of alcohols. Heat mediated antigen retrieval was performed using citrate buffer, pH 6, heated to 95°C for 30 minutes in a water bath followed by washing. Non-specific binding of secondary antibodies was minimised by incubating the tissue sections in a TBS-Tx solution containing 20% normal serum from the species of which the secondary antibodies were raised (Vector Laboratories, Burlingame, CA, USA), for one hour at room temperature, followed by incubation in a cocktail of primary antibodies overnight at 4°C (see SI Table 1 for details of primary antibodies used). The next day, the sections were washed and then incubated at room temperature in a cocktail of appropriate secondary antibodies, conjugated with Alexa Fluor 488, indocarbocyanine (Cy3) or indocarbocyanine (Cy5) (Jackson ImmunoResearch) for one hour. In order to quench autofluorescence, the sections were washed and incubated with an Autofluorescence Eliminator (Millipore) following the manufacturers protocol, for 5 minutes, before being washed and cover-slipped with Vectashield mounting medium (H-1000, Vector Laboratories Inc.).

### Confirmation of antibody specificity

Specificity details for the primary antibodies are provided in SI Table 1. Method specificity was also tested by omitting the primary antibodies in the incubation sequence. To confirm the absence of cross reactivity between IgGs in double and triple immunolabelling experiments, some sections were processed through the same immunohistochemical sequence, except that only an individual primary antibody was applied with the full complement of secondary antibodies.

### Image acquisition

Sections were examined with a confocal laser-scanning microscope (LSM710 or LSM880; Zeiss, Oberkochen, Germany) using a Plan Apochromatic 63 x DIC oil objective (NA 1.4, pixel size 0.13 μm), images were processed using Zen 2009 Light Edition (Zeiss, Oberkochen, Germany) and analysed using ImageJ.

### Colocalisation analyses

The degree of overlap between immunoreactivity for AβO, and citrate synthase and superoxide dismutase 2 (SOD), was quantified to assess the association of AβO with mitochondrial compartments in APP-PSEN mice. The degree of overlap between immunoreactivity for the GABA_A_R α3 subunit and tyrosine hydroxylase (TH), which in LC neurons is located exclusively in cytoplasmic compartments, was quantified to assess the differences in internalised α3-GABA_A_Rs between post mortem control and Alzheimer’s samples. Briefly, images of individual LC neurons were acquired at high resolution. The Coloc2 function in ImageJ was used to quantify the degrees of co-occurrence of overlapping pixels, using the Manders’ overlap coefficient (Manders et al. 1993), as well as the intensities of overlapping pixels, using the Pearson’s correlation coefficient (Aaron et al. 2018). The values for individual cells were grouped and the medians and ranges presented.

### Semi-quantitative analysis of GABA_A_R-α3 immunoreactivity in the LC of tissue from APP-PSEN1 and WT mice

The quantification method was according to our previously published reports (Corteen et al. 2015; Gunn et al. 2013). Using the 100x objective, 10 individual TH immunopositive cells, located within the core region of the LC, were imaged *per* tissue section. The density, area and percentage of the cell region covered by GABA_A_R α3 subunit immunoreactive clusters was quantified using ImageJ (Fiji) and its particle analysis algorithm, using a threshold size of a minimum of 0.04 μm^2^. A single optical section *per* cell, located on the midline of the cell, was used in order to avoid double counting clusters. The dimensions of the optical sections were 84.94 × 84.94 × 0.7 µm in the X, Y and Z planes respectively. This was repeated in n = 4-8 animals (WT and APP-PSEN1 each) which was then replicated in three different litters. All sections were processed and imaged under identical conditions and analyses were performed blind.

### Semi-quantitative analysis of glycine transporter 1 & glycine transporter 2 immunoreactivity in the LC of tissue from APP-PSEN1 and WT mice

In contrast to the uniform distribution of GABA_A_R-α3 subunit immunoreactivity on somata and dendritic domains, labelling for GlyT1 and GlyT2 was largely restricted to the astrocyte plasma membranes and axons respectively. As such, due to the density of astrocytic and axonal processes within the LC, it was not feasible to image individual profiles in any representative manner. Therefore, for these molecules, a larger field of view containing somatic, dendritic or axonally immunopositive processes was used for analyses, rather than individual cells. Nevertheless, the cluster analysis was the same as for GABA_A_R-α3 subunit.

### Semi-quantitative analysis of IBA1 immunopositive cell density in the LC of tissue from APP-PSEN1 and WT mice

Quantification of IBA1-immunopositive cell density was performed according to previously published methods (Seifi et al. 2014). Briefly, TH and IBA1 immunoreactivity within the nuclear core of the LC was imaged in 2-3 tissue sections per animal (n = 10 WT and 10 APP-PSEN1), using a Plan Apochromatic 40 X DIC oil objective (NA 1.3). All IBA1-immunopositive cells within a tissue section were manually counted using ImageJ software (NIH) and mean density (number of cells per unit area) calculated.

### Whole-cell patch clamp electrophysiology recordings in acute brain slices containing the LC

Spontaneous LC firing rate (FR), in the absence and presence of a range of neurotransmitter modulators, was determined in 2-4 month mice, according to previously published protocols (Campos-Lira et al. 2018; Ting et al. 2014). Briefly, mice were killed by cervical dislocation, decapitated and the brain rapidly removed and submerged in room temperature, oxygenated, NMDG-cutting solution, containing (mM): NMDG (92), KCl (2.5), NaH_2_PO_4_ (1.25), NaHCO_3_ (30), Dextrose (25), HEPES (20), sodium ascorbate (5), thiourea (2), sodium pyruvate (3), MgSO_4_ (10), CaCl_2_ (0.5); the pH was adjusted to 7.3 using 12M HCl prior to adding divalent ions. Horizontal 200 μm sections of the brainstem containing the LC were prepared and placed in a holding vial containing pre-warmed 32°C oxygenated NMDG solution for 10 minutes. After this initial recovery period, the slices were transferred to a holding vial containing room temperature, oxygenated, HEPES-holding solution composed of (mM): NaCl (92), KCl (2.5), NaH_2_PO_4_ (1.25), NaHCO_3_ (30), dextrose (25), HEPES (20), sodium ascorbate (5), thiourea (2), sodium pyruvate (3), MgSO_4_ (10), CaCl_2_ (2) for 1 hour at room temperature before recording. A slice was placed in the recording chamber and continuously superfused with oxygenated extracellular solution (ECS) containing (mM): NaCl (126), KCl (2.95), NaH_2_PO_4_ (1.25), NaHCO_3_ (26), dextrose (10), MgSO_4_ (2), CaCl_2_ at 1 ml/min at 32°C.

Recording pipettes contained KMeSO_4_ intracellular solution containing (mM): KMeSO_4_ (135), KCl (5), MgCl_2_ (2), EGTA (0.4), HEPES (10), Mg-ATP (2), Na-GTP (0.5), phosphocreatinine (5) and 0.1 % biocytin (pH 7.3). Recordings were acquired with a Multiclamp 700B amplifier and Digidata 1440A (Molecular Devices) and stored using pClamp 10 software (Molecular Devices). The experimental protocol involved recording baseline cell characteristics in current clamp, including FR (Hz), input resistance (derived from the linear portion of a voltage-current plot of hyperpolarizing current steps MΩ), resting membrane potential (mV), membrane time constant (τ, ms), action potential amplitude (mV), duration (ms) in the presence or absence of bath-applied drugs. Following the recording, the pipette was gently retracted, the slice removed from the recording chamber and submerged in fixative for subsequent neurochemical characterisation of the recorded biocytin-filled neuron (Swinny et al. 2010). Data were analysed with Clampfit software (Molecular Devices).

### Quantitative reverse transcription polymerase chain reaction (qPCR)

qPCR amplifications were performed to assess changes in the mRNA levels for the following inflammatory genes, using pre-designed Taqman primers/probes purchased from Life Technologies (ThermoFisher scientific): IL1β (Mm00434228_m1); TNF-α (Mm00443258_m1); IL 6 (Mm00446190_m1); CSF1 (Mm00432686_m1); CCL2 (Mm00441242_m1) and CCL3 (Mm00441259_g1), in samples from the LC and hippocampus, according to our previously published protocols (Seifi et al. 2018). Gapdh (assay ID: Mm99999915_g1) gene expression was used as the housekeeping gene.

Mice were killed by cervical dislocation and tissue homogenates prepared. RNA was extracted, reverse-transcribed into first-strand cDNA and quantitative PCR (qPCR) amplification was performed in 96-well plates in a Mastermix for probes (Roche, Burgess Hill, UK) and run on a LightCycler® 96 System (Roche). Standard curves were generated for each gene using serial dilutions of a known amount of mRNA extracted from each organ, which were then reverse transcribed into cDNA. Each measurement was performed in duplicate and each Ct value was then converted into ng mRNA using linear regression analysis of the standard curve. Each ng mRNA value was then normalised against the ng housekeeping gene level within the same sample and the mean mRNA levels for every sample was finally calculated and compared across all experimental groups.

### Enzyme-linked immunosorbent assay (ELISA)

An ELISA was used to measure the concentrations of corticosterone (catalogue number MS-E 5000, LDN, Germany) and NA (catalogue number MS-E 5000, LDN, Germany) in blood and brain respectively. Briefly, for corticosterone measurements, mice were killed with an overdose of CO_2_ and blood extracted by cardiac puncture and processed further according to the manufacturer’s protocols. For NA measurements, mice were killed by cervical dislocation and their brains rapidly removed. Two millimetre thick tissue sections containing the LC and hippocampus were obtained using a brain matrix (Harvard Apparatus, #726233). A tissue punch was used to extract the LC and hippocampus regions which were snap frozen using liquid nitrogen until processed further according to manufacturer’s protocols.

### Statistical analysis

The individual statistical tests used to assess specific data sets are indicated in the Results section. An alpha level of less than 0.05 was used to determine statistical significance. For graphical presentation of quantitative data, symbols represent individual data points, bars represent the means and the error bars the SEM. In all cases, GraphPad Prism was used for statistical analyses and graphical presentation of the data.

### Data availability

All data will be deposited and made publicly available in the University of Portsmouth Research Portal (Pure).

## Results

### Expression of AβO within the LC of Alzheimer’s patients and APP-PSEN1 mice

While plaques composed of fibrillar Aβ have been reported in the LC of Alzheimer’s patients, localisation of oligomers in this brain region remain to be demonstrated. Immunoreactivity for an antibody (4G8) that recognises both fibrillar and oligomeric forms of Aβ presented predominantly as extracellular plaques within the LC of post mortem samples from Alzheimer patients, with minimal apparent association with noradrenergic profiles (Fig. 1 A1), in agreement with previously published reports (Thal et al. 2002). No specific signal was detected in human control samples (SI Fig. 1 A). In comparison to Alzheimer’s samples, immunoreactivity in the LC of APP-PSEN1 mice invariably presented as cytoplasmic signal within noradrenergic and non-noradrenergic neurons, with no evidence of extracellular plaques, irrespective of the age of the animal (Fig. 1 A2-4). No specific signal was detected in WT samples (SI Fig. 1 B). In contrast to the LC, cortical regions exhibited an age-dependent expression of Aβ plaques in APP-PSEN1 mice (SI Fig 2A-C), in agreement with previous evidence (Borchelt et al. 1997). In addition, and in contrast to the diffuse cytoplasmic signal detected throughout LC neurons, intracellular Aβ signal in cortical principal neurons was located in close proximity to plasma membranes (SI Fig. 2 D). This suggests that in APP-PSEN1 mice, oligomeric and fibrillar Aβ expression is brain region dependent. Furthermore, since Aβ expression within the APP-PSEN1 LC is primarily soluble and intraneuronal, and thus most likely oligomeric variants, this represents a suitable model to investigate the impact of increased AβO expression on LC function.

**Figure 1.**
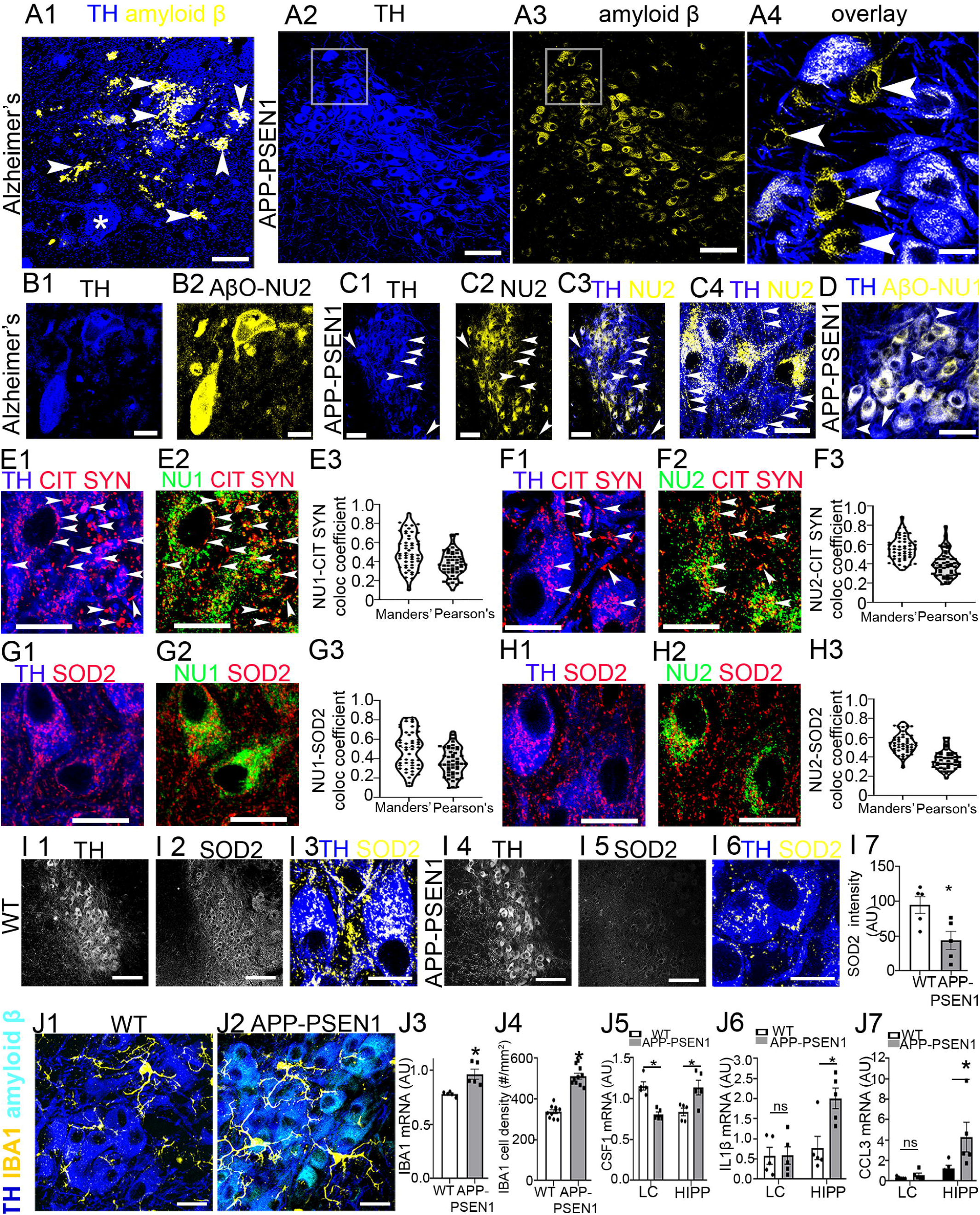
Immunolocalisation of oligomeric amyloid β (Aβ) within the LC of Alzheimer’s patients and APP-PSEN1 mice. (A1) in post mortem samples from Alzheimer’s patients, immunoreactivity for an antibody that recognises both fibrillar and oligomeric forms of Aβ presents exclusively as extracellular plaques (arrowheads) with no intracellular signal evident in LC noradrenergic neurons, identified by immunoreactivity for tyrosine hydroxylase (TH) (asterisk). (A2-3) in APP-PSEN1 mice, immunoreactivity for Aβ is concentrated within LC noradrenergic neurons, with no evidence of extracellular plaques. (A4) is a magnified view of the boxed area in (A 2-3) demonstrating that Aβ is also expressed in non-noradrenergic LC neurons (arrowheads). (B) in LC samples from Alzheimer’s patients, immunoreactivity for an antibody specific for Aβ oligomers (AβO), termed AβO-NU2, shows enrichments within TH-immunopositive somata, together with extracellular clusters. (C1-3) in LC samples from APP-PSEN1 mice, an overview of immunoreactivity for AβO-NU2 reveals its enrichment throughout LC neurons, mirroring the expression pattern in Alzheimer’s samples. However, some TH-immunopositive cells do not exhibit immunoreactivity for AβO-NU2 (arrowheads). (C4) is a magnified view demonstrating that AβO-NU2 immunoreactivity is located within somata of LC neurons and profiles resembling axons (arrowheads). Individual extracellular clusters are also evident. (D) immunoreactivity for a different AβO antibody, termed NU1 that recognises oligomers different in size to NU2, in the LC of APP-PSEN1 mice. Note the significant variation in the intensity of the signal across different TH-immunopositive somata, with some cells (arrowheads) apparently devoid of any immunoreactivity. (E1-2) cytoplasmic AβO-NU1 signal is closely associated (arrowheads) with mitochondria, identified by immunoreactivity for the mitochondria matrix marker citrate synthase. A significant degree of overlap is evident in extracellular clusters which could represent axon terminals. (E3) there is a positive association between the location of immunoreactivity for AβO-NU1 and citrate synthase, with Manders’ representing the degree of co-occurrence of overlapping pixels and Pearson’s the correlation of the intensity of overlapping pixels. (F) in the LC of APP-PSEN1 mice, cytoplasmic AβO-NU2 signal is also closely associated with mitochondria. (G) and (H), in the LC of APP-PSEN1 mice, cytoplasmic immunoreactivity for both AβO-NU1 and AβO-NU2 is closely associated with signal for superoxide dismutase 2 (SOD2), a key mitochondrial enzyme in oxidative stress. (I) in the LC of APP-PSEN1 mice, the level of SOD2 immunoreactivity is significantly decreased. (J1-3) increased expression of the activated microglial marker IBA1 in the LC of APP-PSEN1 mice. Two-tailed unpaired Student’s *t* test. (J4-7) comparative LC and hippocampal mRNA expression levels of the cytokines colony stimulating factor 1 (CSF1), interleukin 1 β (IL1β) and chemokine (C-C motif) ligand 3 (CCL3) respectively. Two way (genotype and brain region) ANOVA. The symbols represent the individual data points, the bars represent the means and the error bars the SEM. * = *P* < 0.05. Scale bars: (A1) 30 µm; (A2-3) 50 µm; (A4) 15 µm; (B1-2) 15 µm; (C1-3) 100 µm; (C4) 15 µm; (D) 60 µm; (E, F, G, H 1-2) 15 µm; (I 1-2, 4-5) 100 µm; (I 3, 6) 15 µm; (J) 15 µm.

To confirm the expression of AβO within LC of samples from Alzheimer’s patients and APP-PSEN1 mice, we used two different antibodies that specifically recognise human AβO in Alzheimer’s, here termed NU1 and NU2 (Lambert et al. 2007). AβO-NU2, produced intense cytoplasmic staining of LC neurons in Alzheimer’s samples, as well as numerous extracellular immunoreactive clusters (Fig. 1 B). No specific signal was detected in human control samples (SI Fig. 3 A). This suggests a hitherto yet described novel significant interaction between LC neurons and the oligomeric forms of Aβ during the disease. An overview of immunoreactivity for AβO-NU2 in the LC of APP-PSEN1 mice revealed a largely intraneuronal pattern that closely mirrored that found in Alzheimer’s samples (Fig. 1 C1-2). Whilst the signal was distributed widely throughout the LC, it was noticeable that some TH-immunopositive cells were devoid of any AβO-NU2 immunoreactivity (Fig. 1 C3). A higher resolution inspection indicated that AβO-NU2 immunoreactivity was concentrated in TH-immunopositive somata, dendrites and fine processes suggestive of axons, and extracellular clusters (Fig. 1 C4). No specific AβO-NU2 signal was detected in WT mouse samples (SI Fig. 3C). Immunoreactivity for AβO-NU1 in APP-PSEN1 mice closely resembled that of AβO-NU2. However, a notable difference was the significant variation in the intensity of signal between different LC TH-immunopositive neurons (Fig. 1 D), suggesting different expression levels across LC noradrenergic neurons. No specific AβO-NU1 signal was detected in WT mouse samples (SI Fig. 3B). We were unable to detect any specific AβO-NU1 in patient samples (data not shown). A possible explanation for the difference between mouse and human expression profiles could be the variation in the expression levels of NU1 in subsets of LC neurons. Since NU1 and NU2 are known to recognise different sized AβO (Lambert et al. 2007), this disparate expression within subsets of noradrenergic neurons suggests that AβO production is not uniform throughout the LC. This AβO-specific parcellation of the LC may underlie the selective vulnerability of subsets of LC principal neurons both in APP-PSEN1 mice (Liu et al. 2008; O’Neil et al. 2007) and Alzheimer’s patients (Chan-Palay and Asan 1989). If AβO-NU1 immunopositive neurons represent a population with vulnerabilities different to NU2-immunopositive subsets, they could have already degenerated in the symptomatic Alzheimer’s patient cohort we drew our samples from. Thus, future studies using samples from patients in the early stages of the condition, which systematically correlate various molecular, morphological and anatomical parameters of the overall LC nucleus with differently sized AβO, will be instrumental in contextualising the role of the Aβ system in LC integrity in Alzheimer’s.

The significant cytoplasmic signal for AβO-NU1/2 raised the question whether AβO are associated with specific organelles. A proposed pathway for intracellular AβO in Alzheimer’s pathogenesis is its incorporation into the mitochondrial matrix and disruption of cellular bioenergetics (Hansson Petersen et al. 2008). Immunoreactivity for citrate synthase, an enzyme located within the mitochondrial matrix (Wiegand and Remington 1986), was used to explore the association between AβO-NU1/2 and mitochondrial compartments in APP-PSEN1 mice. There was a significant overlap between somatic immunoreactivity for AβO-NU1 (Fig. 1 E1-2) and AβO-NU2 (Fig. 1 F1-2) with the signal for citrate synthase. Quantification of the degree of colocalised pixels between citrate synthase:AβO-NU1 (Fig. 1 E3) and citrate synthase:AβO-NU2 (Fig. 1 F3) revealed positive correlations for both the degrees of co-occurrence of overlapping pixels, using the Manders’ overlap coefficient (Manders et al. 1993), [(NU1: median, range; 0.5, 0.1 - 0.9, n = 100 cells, Fig. 1 E3), (NU2: median, range: 0.6, 0.33-0.9, n = 100 cells, Fig. 1 F3) as well as the intensities of overlapping pixels, using the Pearson’s correlation coefficient (Aaron et al. 2018) [(NU1: median, range; 0.4, 0.1 - 0.7, n = 100 cells, Fig. 1 E3), (NU2: median, range: 0.4, 0.2-0.8, n = 100 cells, Fig. 1 F3). There was also a striking overlap of citrate synthase-AβO-NU1/2 for immunoreactive clusters neighbouring the periphery of somata and dendrites, which could be indicative of the locations of axonal terminals. This indicates that in LC neurons of APP-PSEN1 mice, intraneuronal AβO are located in close proximity to mitochondria.

A consequence of AβO-mediated mitochondrial dysfunction is oxidative stress (De Felice et al. 2007; Picone et al. 2014). Importantly, mitochondrial oxidative stress is directly associated with LC FR, and negatively affects neuronal viability leading to neurodegeneration (Sanchez-Padilla et al. 2014). A key mitochondrial enzyme in this process is superoxide dismutase 2 (SOD2) which provides protection against AβO-mediated oxidative stress (Du et al. 2019) and its expression is decreased in Alzheimer’s (Majd and Power 2018). We therefore assessed the association of AβO-NU1/2 expression with that of SOD2 in APP-PSEN1 mice. We detected a positive association for the location of SOD2 immunoreactive clusters with both AβO-NU1 (Fig. 1 G) and AβO-NU2 (Fig. 1 H). In comparison to WT samples, there was a noticeable decrease in the intensity of SOD2 immunoreactivity in the LC of APP-PSEN1 mice (Fig. 1 I1-6), which was borne out by a quantitative assessment of the respective fluorescence intensities, in samples reacted and imaged under identical conditions (*P* = 0.02, two-tailed unpaired Student’s t test; n = 5 animals) (Fig. 1 I7). This collectively suggests a strong association of intraneuronal AβO with mitochondrial compartments of LC neurons and alterations in cellular bioenergetics regulatory mechanisms, which have the potential to adversely impact on their long-term viability.

Neuroinflammation is considered a core component of Alzheimer’s pathology with both Aβ fibrils (Heneka et al. 2015) and AβO mediating specific pathways (Welikovitch et al. 2020). A distinctive feature of Alzheimer’s related neuroinflammation is the presence of activated microglia. We therefore assessed the LC of APP-PSEN1 mice for signs of altered immune status, using the microglia/macrophage-specific calcium-binding protein ionized calcium binding adaptor molecule 1 (IBA1), the expression of which is increased in Alzheimer’s-associated neuroinflammation (Heneka et al. 2015). TH-immunopositive neurons in the LC of APP-PSEN1 mice appeared to be contacted by a greater number of IBA1-immunoreactive profiles which were also more ramified, compared to WT, suggesting increased expression levels of this inflammatory marker (Fig. 1 J1-2). This increased expression was confirmed at both mRNA (*P* = 0.0053, two-tailed unpaired Student’s t test; n = 10 animals) (Fig. 1 J3) and cell density levels (*P* <0.0001, two-tailed unpaired Student’s t test; n = 10 animals) (Fig. 1 J4). The production of inflammatory cytokines is considered to be a consequence of microglial activation in Alzheimer’s (Heneka et al. 2015). Since the LC of APP-PSEN1 mice exhibits only the oligomeric forms of Aβ, whilst oligomeric as well as fibrillar expression is evident in cortical regions, we assessed whether there were differences in the levels of inflammatory cytokines between the LC and hippocampus. Accordingly, we detected contrasting cytokine expression profiles in the LC and hippocampus. mRNA expression for colony stimulating factor 1 receptor (CSF1), which is anti-inflammatory and prevents the progression of Alzheimer’s-like pathology (Olmos-Alonso et al. 2016), was significantly decreased in the LC (*P* = 0.0013, 2-ANOVA, Sidak’s; n = 5 animals), yet upregulated in the hippocampus (*P* = 0.0045, 2-ANOVA, Sidak’s; n = 5 animals) (Fig. 1 J5). Since CSF1 signalling purportedly promotes neuroprotection (Luo et al. 2013), its contrasting expression profiles in the LC and hippocampus, as a result of Alzheimer’s-like pathology, could be a contributing factor to the early vulnerability of the LC to Alzheimer’s. In contrast, whilst the mRNA levels for interleukin 1 beta (IL-1β) (*P*= 0.0081, two-way ANOVA with Sidak’s; n = 5 animals) (Fig. 1 J6) and chemokine (C-C motif) ligand 3 (CCL3) (*P* = 0.0216, 2-ANOVA, Sidak’s; n = 5 animals) (Fig. 1 J7) were significantly increased in hippocampal samples, no significant changes were detected in the LC. We found no significant differences in the mRNA expression, in both LC and hippocampus, for TNFα, IL6 and CCL2 (data not shown). This indicates that in this model, the LC inflammatory cascades are distinct to cortical domains. This may signal a unique disease phenotype for this brain region, as a result of Aβ expression occurring exclusively in the oligomeric form, compared to cortical regions which include the fibrillar variants as well.

### AβO is associated with LC synapses and induces LC neuronal hyperexcitability in APP-PSEN1 mice

Apart from the cytoplasmic expression profile of AβO-NU1-2 outlined above, numerous extracellular immunoreactive clusters that decorated LC neuronal surfaces were also evident. As outlined previously, AβO have been shown to interact at synaptic domains. Furthermore, AβO-NU1-2, in hippocampal samples, have previously been shown to exhibit a punctate dendritic labelling pattern which may be synonymous with synaptic junctions (Lambert et al. 2007). Therefore, we next assessed the association of AβO immunoreactivity with signal for synaptic marker proteins. In APP-PSEN1 mice, clusters immunoreactive for AβO-NU2 were located in close apposition to puncta immunopositive for the synaptic vesicle glycoprotein 2A (SV2A) (Fig. 2 A), a protein expressed in excitatory and inhibitory synapses and is thus representative of the location of all synaptic junctions (Bajjalieh et al. 1994). Furthermore, AβO-NU2 immunoreactivity was in close contact with clusters immunoreactive for the vesicular GABA transporter (VGAT), a marker of inhibitory synapses (Fig. 2 B) as well as the vesicular glutamate 2 (VGLUT2), a marker of excitatory synapses (Fig. 2 C).

**Figure 2.**
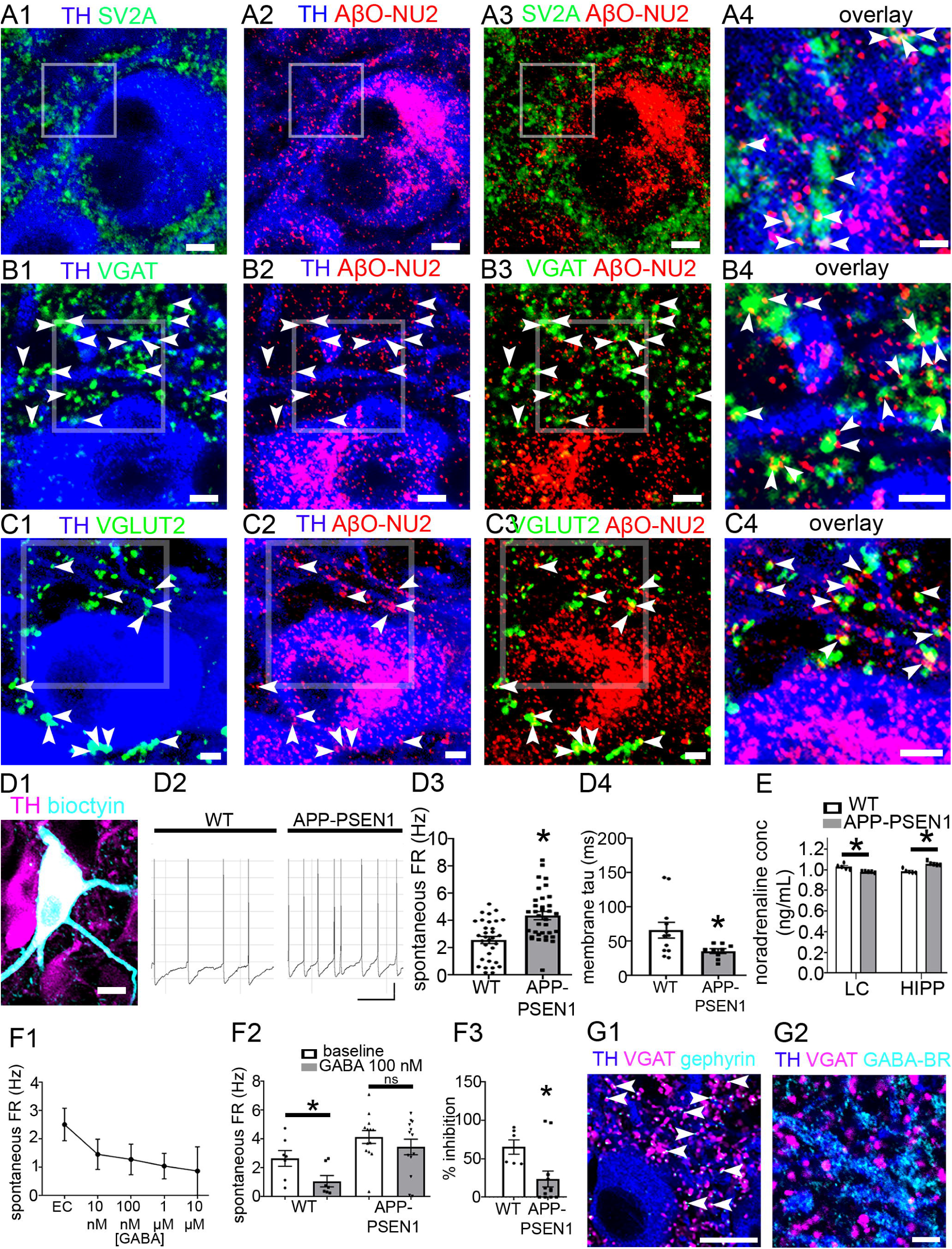
Association of AβO with synapses and LC neuronal excitability in APP-PSEN1. (A) extracellular AβO-NU2 immunoreactivity is located in close apposition to clusters immunopositive for the pan-presynaptic marker protein, synaptic vesicle glycoprotein 2A (SV2A) (arrowheads), indicating expression at synaptic junctions. (A4) is a magnified view of the boxed areas in (A1-3). (B) extracellular AβO-NU2 immunoreactive clusters are closely associated with inhibitory axon terminals, identified by immunoreactivity for the vesicular GABA transporter (VGAT) (arrowheads), indicating expression in close proximity to GABAergic synapses. (B4) is a magnified view of the boxed areas in (B1-3). (C) extracellular AβO-NU2 immunoreactive clusters are closely associated excitatory axon terminals, identified by immunoreactivity for the vesicular glutamate transporter 2 (VGLUT2) (arrowheads), indicating expression in close proximity to glutamatergic synapses. (C4) is a magnified view of the boxed areas in (C1-3). (D1) representative image of a recorded neuron, loaded with biocytin during the recording, and processed post-hoc for TH immunohistochemistry, confirming that recorded neurons were located within the LC and were noradrenergic. (D2) representative traces of the spontaneous firing rates (FRs) of LC noradrenergic neurons in WT and APP-PSEN1 mice. Note the significantly higher FR in the trace for APP-PSEN1. (D3) quantification of the spontaneous FR (Hz) of LC noradrenergic neurons in WT and APP-PSEN1 mice. The data points represent individual cells. Two-tailed unpaired Student’s *t* test. (D4) quantification of the membrane time constant for WT and APP-PSEN1 mice. The data points represent individual cells. Two-tailed unpaired Student’s *t* test. (E) quantification of the concentration of noradrenaline in the LC and hippocampus of WT and APP-PSEN1 mice using an ELISA for NA. The data points represent individual mice. Two way (genotype and brain region) ANOVA. (F1) concentration-dependent effect of the inhibitory role of GABA on spontaneous LC FR in WT mice. (F2) diminished inhibitory effect of GABA in LC neurons form APP-PSEN1 mice. The data points represent individual cells. Two way (genotype and drug) ANOVA. (F3) percentage decrease in spontaneous LC FR following the application of 100 nm GABA in WT and APP-PSEN1 mice. The data points represent individual mice. Two-tailed unpaired Student’s *t* test. (G1) demonstration that the majority of GABA release sites, identified by immunoreactivity for the vesicular GABA transporter (VGAT) are located adjacent to domains containing putative GABA_A_ and glycine receptors, identified by immunoreactivity for the GABA_A_ and glycine receptor anchoring protein gephyrin. (G2) in contrast to (F1), sparse clusters immunoreactive for the GABA_B_ receptor are located on TH-immunopositive profiles which rarely are located adjacent to VGAT-immunoreactive clusters. The symbols represent the individual data points, the bars represent the means and the error bars the SEM. * = *P* < 0.05. Scale bars: (A1-3) 5 µm; (A4) 1 µm; (B1-3) 5 µm; (B4) 3 µm; (C) 2 µm; (D1) 10 µm; (D2) horizontal, 500 ms, vertical 20 mV; (G1) 10 µm; (G2) 5 µm.

Given the important role that synaptic transmission has in setting the spontaneous FR of LC neurons, and the established impact of AβO on synaptic transmission discussed previously, we next assessed the functional consequences of the increased AβO expression in APP-PSEN1 mice, by recording LC spontaneous FRs from identified NA LC neurons (Fig. 2 D). APP-PSEN1 LC neurons exhibited significantly greater FRs (*P* < 0.001; two-tailed unpaired Student’s *t* test; n = 32 WT and 35 APP-PSEN1 cells) (Fig. 2 D2-3) and decreased membrane time constant compared to WT (*P* = 0.038, two-tailed unpaired Student’s t test, n = 9 cells) (Fig. 2 D4). No other changes in LC physiological parameters were detected (Table 1). Accompanying this enhanced LC excitability was a significant decrease in NA concentration in the LC (*P* = 0.0032, 2-ANOVA, n =5 animal), and a significant increase in the hippocampal content (*P* < 0.0001, 2-ANOVA, n =5 animal) (Fig. 2 E).

**Table 1.** Summary of basic membrane characteristics of recorded noradrenergic LC neurons in WT and APP-PSEN1 mice. Data are presented as the mean ± SEM. n = number of cells used per parameter.

| **Parameter**<br>**(N) = number of cells** | **WT** | **APP-PSEN1** | **P (unpaired Student's *t* test)** |
| --- | --- | --- | --- |
| **Spontaneous FR, Hz (n)** | 2.57 ± 0.26 (32) | 4.37 ± 0.30 (35) | < 0.0001 |
| **Resting membrane potential, mV (n)** | -53.20 ± 1.23 (12) | -52.76 ± 1.29 (9) | 0.8083 |
| **Cell capacitance, pF (N)** | 30.45 ± 1.25 | 30.76 ± 1.86 | 0.8884 |
| **Threshold for action potential, mV (n)** | -43.27 ± 1.71 (11) | -44.33 ± 2.49 (9) | 0.7225 |
| **Amplitude of action potential, mV (n)** | 62.82 ± 2.39 (11) | 66.11 ± 3.05 (9) | 0.3997 |
| **Cells spontaneously firing action potentials, % (n)** | 94.12 (34) | 97.22 (36) | NA |
| **Input resistance, MΩ (n)** | 386.6 ± 0.05 (12) | 295.4 ± 0.04 (8) | 0.1840 |
| **Time constant, ms (n)** | 66.3 ± 11.45 (12) | 35.88 ± 3.48 (9) | 0.0380 |

In light of this heightened excitability of LC neurons, we next assessed whether this could be due to changes in inhibitory modulation. GABA and glycine provide the predominant inhibitory modulation of LC neurons (Somogyi and Llewellyn-Smith 2001). We therefore explored whether the increased levels of AβO in AP-PSEN1 impacts on these pathways in regulating basal spontaneous FR. Using a recording electrode solution that is representative of the native intracellular Cl^-^ ion concentration, bath applied GABA induced a concentration dependent decrease in the spontaneous FR of WT LC neurons with an approximate IC_50_ of 100 nM (Fig. 2 F1). With this concentration of GABA, there was a significant drug [F _(1, 18)_ = 12.38, *P* = 0.0025, two-way ANOVA with repeated measures (2-RMA)] and genotype [F _(1, 18)_ = 7.685, *P* = 0.0126, 2-RMA] effect on FR. However, *post-hoc* analysis revealed that while GABA (100 nM) significantly inhibited FR in WT cells (*P* = 0.0142, Sidak’s; n = 7 cells), this effect was abolished in cells from APP-PSEN1 mice (*P* = 0.1665, Sidak’s; n = 12 cells) (Fig. 2 F2). Calculating the percentage of inhibition induced by this concentration of GABA revealed a significant decrease in response of APP-PSEN1 neurons (*P* = 0.0201, unpaired Student’s t test) (Fig. 2 F3). This impaired GABAergic receptor response could be due to changes in either GABA_A_ or GABA_B_ receptor (GABA_A-B_R) subtypes. However, a previous electrophysiological report demonstrated that the predominant effect of GABA on LC neurons was mediated *via* GABA_A_Rs, with only significantly higher concentrations (> 600 µM) activating GABA_B_Rs (Osmanovic and Shefner 1990). We verified this predominant association of GABA with GABA_A_Rs at the molecular level with immunoreactive clusters for the vesicular GABA transporter (VGAT), a marker of GABA and glycine release sites, being immediately adjacent to cluster immunoreactive for gephyrin, a protein that anchors GABA_A_R and GlyRs in postsynaptic junctions (Fig. 2 G1). In contrast, sparse immunoreactivity for GABA_B_R was detectable in the LC, which was infrequently associated with VGAT immunoreactivity (Fig. 2 G2). We therefore focussed further on GABA_A_Rs and GlyRs.

### Impaired α3-GABA_A_Rs in LC neurons of Alzheimer’s patients and APP-PSEN1 mice

Complex changes in the native expression patterns of various GABA_A_R subunits have been reported in a range of cortical brain regions of Alzheimer’s patients (Kwakowsky et al. 2018). While the complement of GABA_A_R subunits expressed within the mouse and human LC nucleus has been demonstrated (Waldvogel et al. 2010), confirmation of the neurochemical identity of the LC cell types expressing specific subunits, at high resolution, remains to be reported. Therefore, before we were able to identify any changes in samples from Alzheimer’s patents and APP-PSEN1 mice, it was imperative to describe the native expression patterns, using WT mice and control human tissue. In agreement with a previous report in rat (Corteen et al. 2011), the GABA_A_R α1 subunit was exclusively expressed in TH immunonegative cells of the mouse (Fig. 3 A1-3). The restriction of this subunit to non-NA neurons of the LC was confirmed in human samples as well (Fig. 3 A4). In contrast, immunoreactivity for the α3 subunit in this region of the brainstem strictly overlapped with that of TH, both within the LC nuclear core and the pericoeruleur dendritic regions of the mouse LC (Fig. 3 B1-2), suggesting a significant contribution of this subunit to GABA-mediated regulation of the LC-NA system. The specificity of this labelling pattern was confirmed using tissue from α3 subunit gene deleted mice (α3^-/-^) in which no specific signal was detectable (Fig. 3 B3-4).

**Figure 3.**
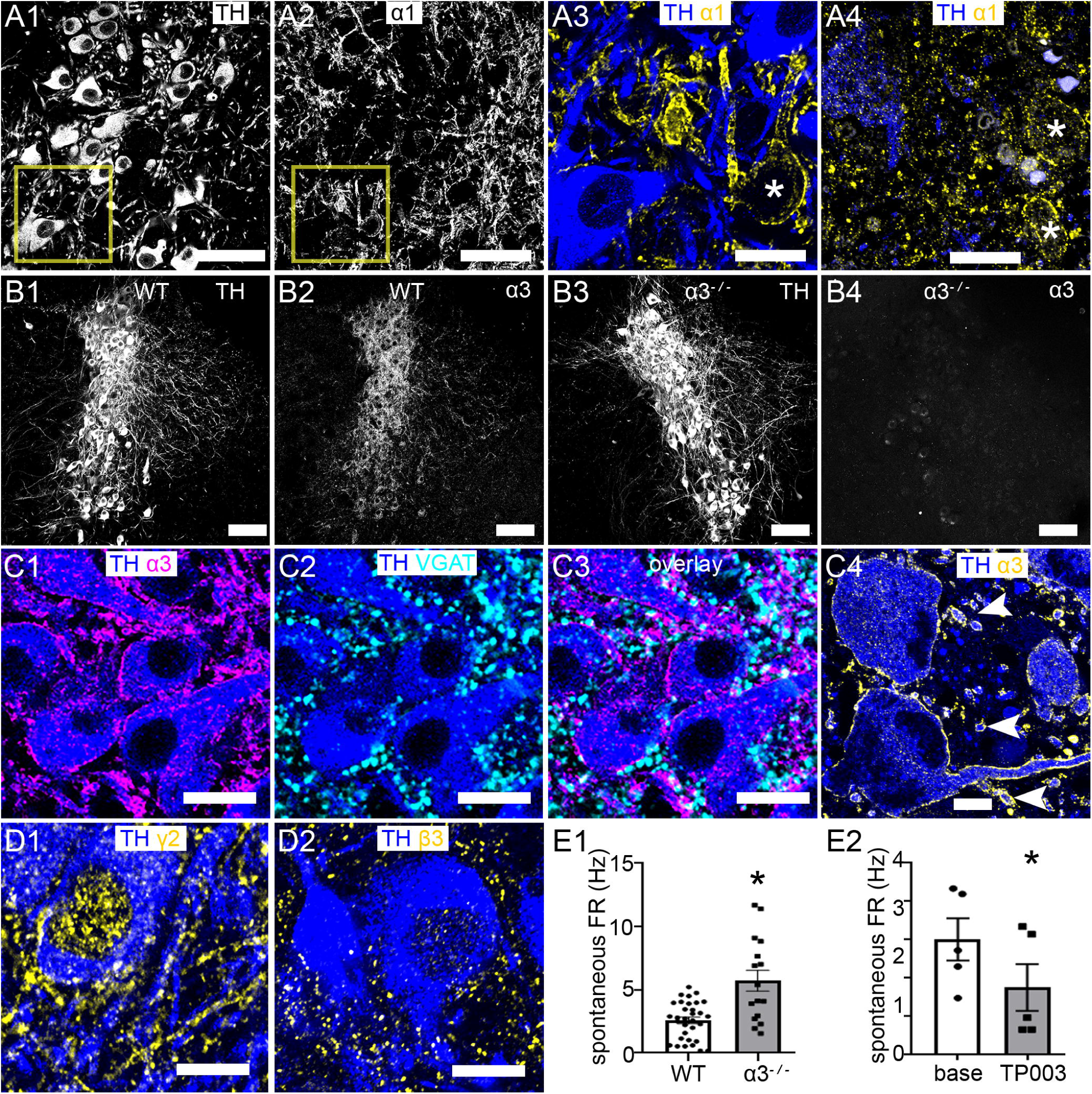
Demonstration of native GABA_A_ receptor (GABA_A_R) subtypes expressed in mouse and human LC neurons. (A) the GABA_A_R α1 subunit is expressed in non-noradrenergic neurons (asterisks) of the (A1-3) mouse and (A4) human LC. (B) overview of LC α3 subunit immunoreactivity in WT and α3 gene deleted mice (α3 ^-/-^), thereby confirming the specificity of the α3 antibody used. (C1-3) high resolution immunolocalisation demonstrating that in the mouse LC, α3 immunoreactivity is concentrated on somatic and dendritic surfaces adjacent to GABA release sites, identified by immunoreactivity for VGAT. (C4), in the human LC, α3 immunoreactivity is also enriched on the somatic and dendritic (arrowheads) of noradrenergic neurons. Mouse LC noradrenergic neurons also express (D1) the γ2 and (D2) β3 subunits. (E) α3-containing GABA_A_Rs (α3-GABA_A_Rs) contribute to the basal FR of LC neurons, with (E1) the deletion of this subunit resulting in a significant increase (two-tailed unpaired Student’s t test), whilst (E2) its pharmacological activation resulting in a significant decrease (two-tailed unpaired Student’s *t* test). The symbols represent the individual data points, the bars represent the means and the error bars the SEM. * = *P* < 0.05. Scale bars: (A1-2) 30 µm; (A3) 10 µm; (A4) 20 µm; (B) 100 µm; (C) 15 µm; (D) 10 µm.

High resolution inspection revealed that the majority of α3 subunit immunoreactive clusters were located on plasma membrane surfaces of TH immunopositive neurons immediately adjacent to GABA release sites, identified by immunoreactivity for VGAT in mouse (Fig. 3 C1-3). This pattern of α3 subunit immunoreactivity being concentrated on somatodendritic surfaces of LC NA neurons was replicated in control human samples (Fig. 3 C4). We were unable to detect any specific signal for the α2 subunit (or any other α subunit) in LC neurons, even though we were able to reproduce the quintessential immunoreactivity pattern for this subunit which others have demonstrated in other brain regions such as the hippocampus (SI Fig. 4). TH immunopositive neurons also displayed immunoreactivity for the γ2 (Fig. 3 D1) and β3 (Fig. 3 D2) subunits, suggesting that α3-β3-γ2 are the subunit combinations forming the major GABA_A_R subtype on LC NA neurons.

The importance of α3-GABA_A_Rs in setting the spontaneous LC FR was confirmed in cells from α3^-/-^ mice, which displayed a significantly higher basal FR compared to WT littermates (*P* = 0.0048, two-tailed unpaired Student’s *t* test; n = 21 WT cells and 16 α3^-/-^ cells) (Fig. 3 E1). Additionally, the α1/2/3/5-GABA_A_Rs positive allosteric modulator TP003 (Neumann et al. 2018), which is likely to induce its effects on LC-NA neurons predominantly via α3-β/γ-GABA_A_Rs due to the absence of other α subunits in this cell type, significantly inhibited the FR of neurons from WT mice (*P* = 0.0099, two-tailed paired Student’s *t* test; n = 5 cells) (Fig. 3 E2). Presumably this effect of TP003 reflects an action of the drug to enhance the effects of ambient GABA on α3-GABA_A_Rs. Collectively, these data suggest that α3-GABA_A_Rs are integral in setting the basal level of LC neuronal activity, and therefore LC-mediated brain functions and behaviours.

Given the above-mentioned importance of α3-GABA_A_Rs to LC activity, we next explored their expression and function in APP-PSEN1 mice. Clusters immunoreactive for the α3 subunit were noticeably fewer and smaller in APP-PSEN1 samples, with the majority of the signal located within the cytoplasm of TH immunopositive neurons, rather than the characteristic clustering on plasma membrane surfaces (Fig. 4 A, B). We detected significant decreases in α3 cluster density (*P* < 0.0001, two-tailed unpaired Student’s *t* test; n = 40 cells from 4 WT animals and n = 50 cells from 5 APP-PSEN1 animals), size (*P* < 0.0001; two-tailed unpaired Student’s *t* test), fluorescence intensity (*P* < 0.0001; two-tailed unpaired Student’s *t* test) and percentage of the cell area they covered (*P* < 0.0001; two-tailed unpaired Student’s *t* test) (Fig. 4 C). In APP-PSEN1 mice, there was a noticeable overlap between pools of α3 and AβO-NU2 immunopositive clusters, both within the cytoplasm of somata and dendrites (Fig. 4D). The functional consequences of this Aβ-altered α3 profile was confirmed by TP003 which significantly decreased the FR of WT cells (*P* = 0.04, 2-RMA, Sidak’s; n = 5 cells), yet had a negligible and non-significant effect in APP-PSEN1 cells (*P* = 0.9894, 2-RMA, Sidak’s; n = 6 cells) (Fig. 4 E).

**Figure 4.**
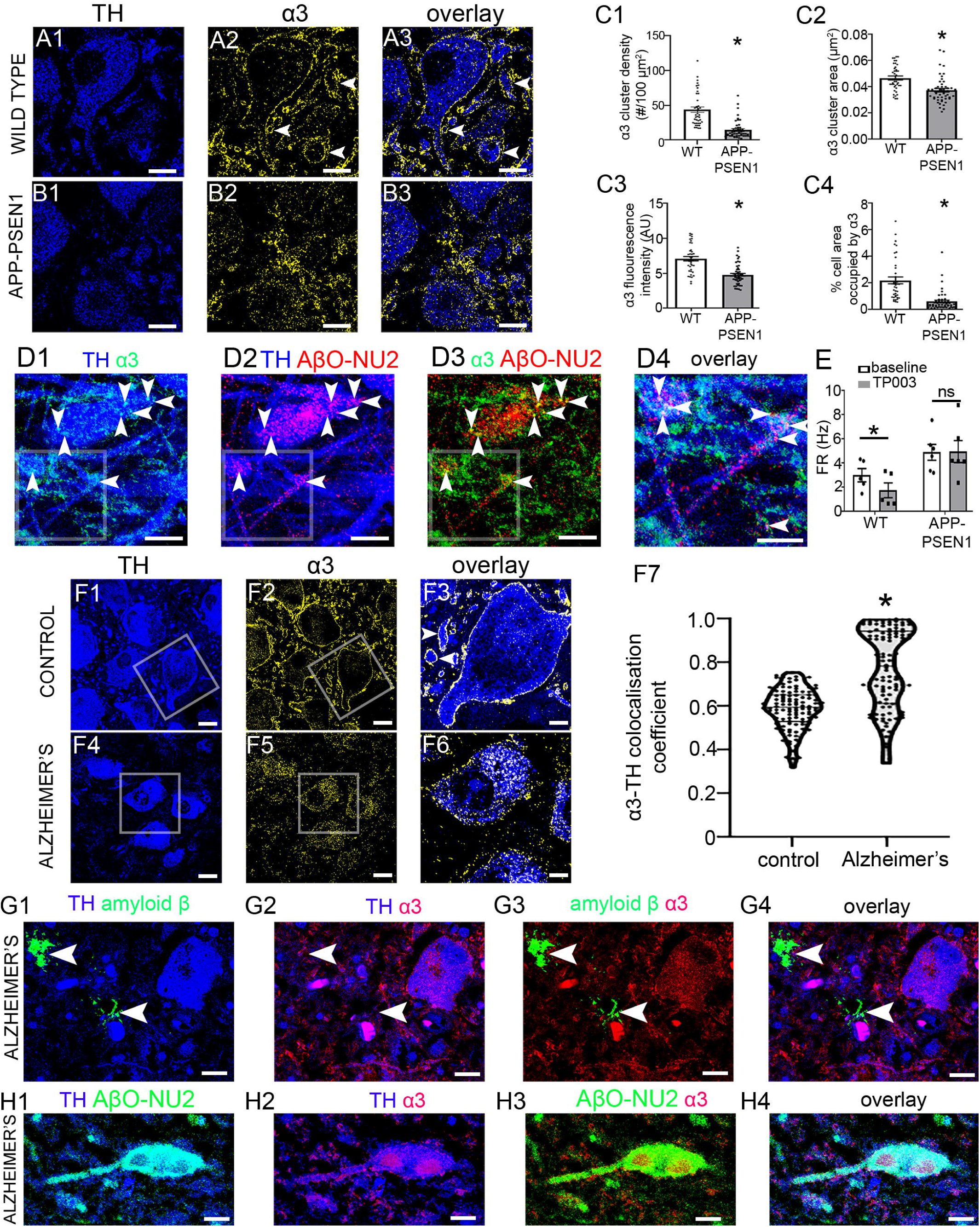
Aβ and α3-GABA_A_Rs in the LC of Alzheimer’s patients and APP-PSEN1 mice. (A) the immunoreactivity pattern of the α3 subunit in the LC of a WT mouse. (B) the immunoreactivity pattern of the α3 subunit in the LC of an APP-PSEN1 mouse, processed and imaged under conditions identical to WT samples. Note the sparse localisation of signal on somato-dendritic plasma membranes, with the majority of the signal preferentially located in cytoplasmic compartments. Quantification of the (C1) density, (C2) size, (C3) mean fluorescence intensity of α3 subunit immunoreactive clusters and (C4) percentage of cell area they occupy, in the LC of WT and APP-PSEN1 mice. Two-tailed unpaired Student’s t test. (C) shows that in LC somata and dendrites of APP-PSEN1 mice, pools of AβO-NU2 immunopositive clusters overlap with clusters immunopositive the α3 subunit (arrowheads). (D4) is a magnified view of the boxed areas in (D1-3). (D) differences in the inhibitory effect of the α1/3/5-GABA_A_R positive allosteric modulator TP003 in LC neurons from WT and APP-PSEN1 mice. While TP003 robustly inhibits LC FR in WT cells, there is no significant effect in cells from APP-PSEN1 mice. The data points represent individual cells. Two-way (genotype and drug) ANOVA. (F1-3) immunoreactivity for the α3 subunit in the LC of control human tissue. (F4-6) immunoreactivity for the α3 subunit in the LC of an Alzheimer’s patient. Note how the α3 signal in patient material mirrors that of the APP-PSEN1 mouse, being predominantly restricted to cytoplasmic compartments, with significantly lower levels evident on plasma membrane compartments. (F7) quantification of the association between immunoreactivity for the α3 subunit and TH, as a measure of cytoplasmically located α3 subunit, using the Manders’ coefficient of the co-occurrence of overlapping pixels. The data points represent individual cells. P > 0.05, Kolmogorov–Smirnov test, n = 110 cells per group. (G) shows the lack of an association between signal for fibrillar Aβ (arrowheads) and the α3 subunit in the LC of Alzheimer’s patients. (H) In the LC of Alzheimer’s patients, immunoreactivity for AβO-NU2 and the α3 subunit are located in close proximity to one another, in a pattern that mirrors expression in APP-PSEN1 mice. The symbols represent the individual data points, the bars represent the means and the error bars the SEM. * = *P* < 0.05. Scale bars: (A, B, D1-3) 10 µm; (D4) 5 µm; (F1-2; F4-5) 15 µm; (F3, F6) 6 µm; (G) 15 µm; (H) 20 µm.

We sought to confirm these AβO-mediated changes in LC α3 subunit expression using post mortem LC samples from Alzheimer’s patients with demonstrated Aβ expression in this brain region. The α3 subunit immunoreactivity pattern in such samples was indistinguishable to that obtained in APP-PSEN1 mice, being preferentially located in cytoplasmic compartments occupied by TH immunoreactivity (Fig. 4 F1-6). This preponderance of α3 expression in cytoplasmic rather than membrane domains in Alzheimer’s tissue was confirmed by quantifying the degree of colocalisation between immunoreactivity for α3 and TH, since TH is expressed exclusively in cytoplasmic compartments. There was a significant difference in the distributions for the α3-TH Manders’ colocalisation coefficients between control and Alzheimer’s samples (*P* <0.0001, Kolmogorov-Smirnov, n = 110 cells from 5 patients in each category) (Fig. 4 F7). In such patients’ samples, there was no discernible interaction between immunoreactivity for α3 and Aβ immunopositive plaques (Fig. 4 G). In stark contrast, a significant association between α3 and AβO expression was evident with the considerable overlap between these signals throughout neuronal compartments (Fig. 4 H). These data collectively suggest that in Alzheimer’s there is a significant AβO-impairment in a key regulator of LC excitability, which in turn could have a far-reaching impact on the functioning of the LC-NA system throughout the brain.

### GlyRs and transporters are resilient to AβO pathology in APP-PSEN1 mice and their modulation reverses LC hyperexcitability

We next explored whether the inhibitory drive arising from the LC GlyR-GlyT systems is also influenced by the elevated levels of AβO in APP-PSEN1 mice, firstly by characterising these systems in the WT mouse. Immunoreactivity for GlyR subunits was located throughout somatodendritic surfaces of TH immunopositive neurons, in close proximity to VGAT immunopositive clusters (Fig. 5 A), and their pharmacological activation decreased LC FR (Fig. 5 B). Immunoreactivity for the glycine transporters one and two (GlyT1-2) (Eulenburg et al. 2005) was concentrated on astrocytic and axonal profiles respectively (Fig. 5 C1-3, D1-3). In addition, inhibition of GlyT1, using ASP2535 (Harada et al. 2012) (1 µM) (*P* = 0.0015, two-tailed paired Student’s t test; n = 8 cells) and GlyT2, using ALX1393 (Eckle and Antkowiak 2013)(1 µM), (*P* = 0.0006; two-tailed paired Student’s t test; n = 11 cells) both significantly decreased LC FR (Fig. 5 C4, D4). Thus, the activity of both GlyRs and GlyT subtypes contributes to setting the basal LC FR.

**Figure 5.**
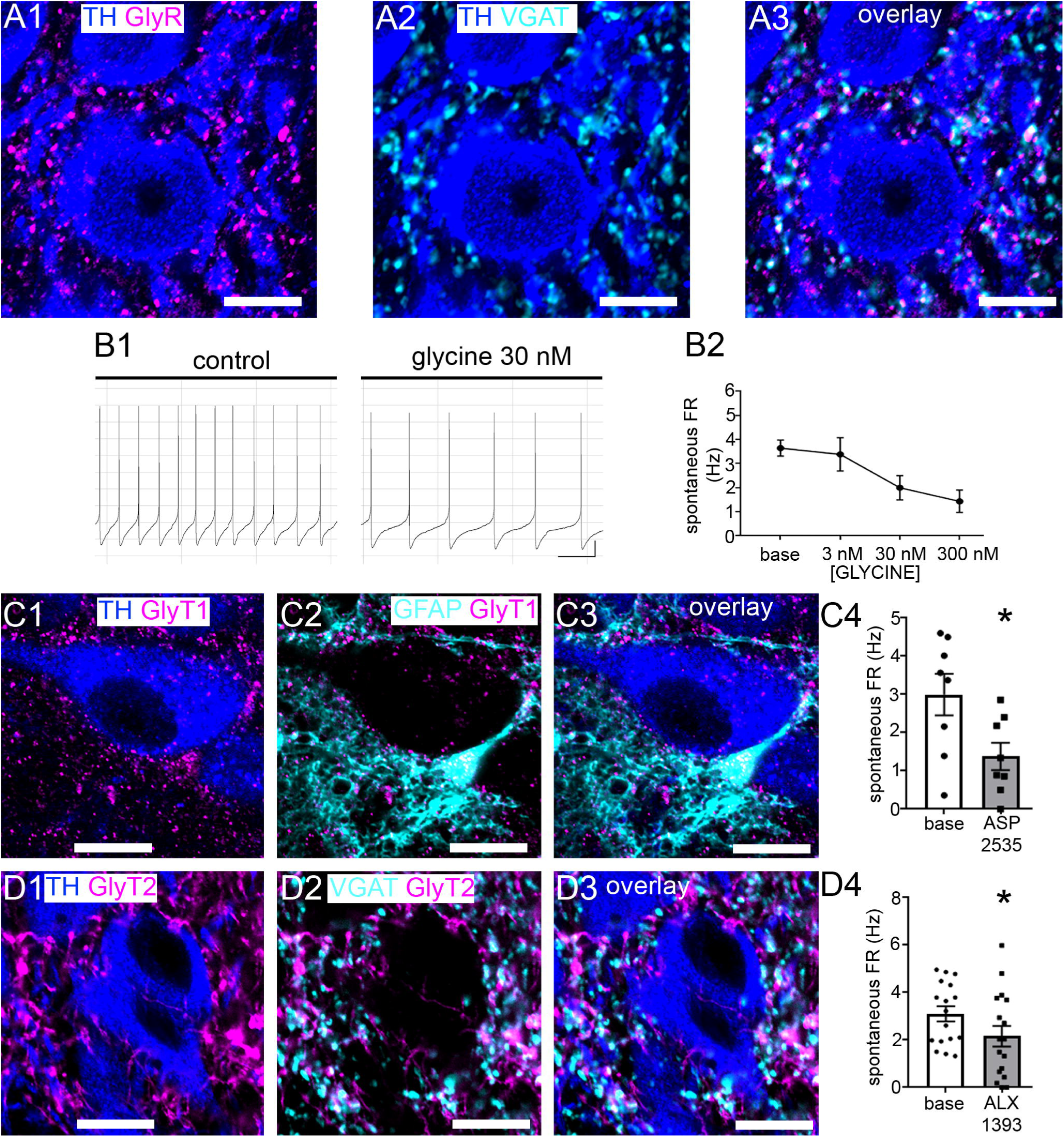
Demonstration of native glycine receptor and transporter subtypes expressed in mouse LC neurons. (A) immunoreactivity for an antibody that recognises all GlyR α subunits and its close association with immunoreactivity for the vesicular GABA transporter (VGAT). (B) concentration-dependent inhibitory effect of applied glycine on spontaneous LC FR. (C1-3) immunoreactivity for the glycine transporter 1 (GlyT1) is associated with astrocytic profiles, identified by immunoreactivity for glial fibrillary acid protein (GFAP). (C4) quantification of the frequency (Hz) of LC noradrenergic neuronal spontaneous firing before and after the bath application of 1 μM ASP-2535, an inhibitor of GlyT1. Two-tailed unpaired Student’s *t* test. (D1-3) immunoreactivity for the glycine transporter 2 (GlyT2) is associated with inhibitory axon terminals, identified by the immunoreactivity for VGAT. (D4) quantification of the frequency (Hz) of LC noradrenergic spontaneous firing before and after the bath application of 1 μM ALX-1393, an inhibitor of GlyT2. Two-tailed unpaired Student’s *t* test. The symbols represent the individual data points, the bars represent the means and the error bars the SEM. * = *P* < 0.05. Scale bars: (A) 10 µm; (C, D) 20 µm.

In light of the above potential to pharmacologically exploit this coexisting inhibitory system as a means to regulating LC excitability, we next characterised the GlyR-GlyT systems in the APP-PSEN1 model. There were no differences between WT and APP-PSEN1 mice in terms of the pattern (Fig. 6 A1-4), density (*P* = 0.5299, two-tailed unpaired Student’s t test; n = 8 animals) (Fig. 6 A5) and cluster size (*P* = 0.9250, two-tailed unpaired Student’s t test; n = 8 animals) (Fig. 6 A6) of GlyR immunoreactive clusters. Importantly, glycine (30 nM) significantly reduced APP-PSEN1 LC neurons FRs [F _(1, 17)_ = 24.05, *P* = 0.0001; 2-RMA] in LC FR for both WT (*p* = 0.0140, Sidak’s; n = 10) and APP-PSEN1 (*P* = 0.0026, Sidak’s; n = 9) cells (Fig. 6 A7). Although there was a decrease in GlyT2 immunoreactive cluster density (*P* = 0.0341, two-tailed unpaired Student’s t test; n = 7 animals) (Fig. 6 B1-5) and cluster size (*P* = 0.0244, two-tailed unpaired Student’s t test; n = 7 animals) (Fig. 6 B6), no such changes were detected for GlyT1 (data not shown). This was further confirmed, using the GlyT1 inhibitory ASP2535, which also reversed LC hyperexcitability in LC neurons [F (1, 14) = 54.77, *P* < 0.0001, 2-RMA) of WT (*P* = 0.0006, Sidak’s; n = 10 cells) and APP-PSEN1 (*P* = 0.0001, Sidak’s; n = 10 cells) mice (Fig. 6 B7). This therefore identifies this pathway as a potential strategy for reversing aberrant levels of LC excitability due to Alzheimer’s pathology.

**Figure 6.**
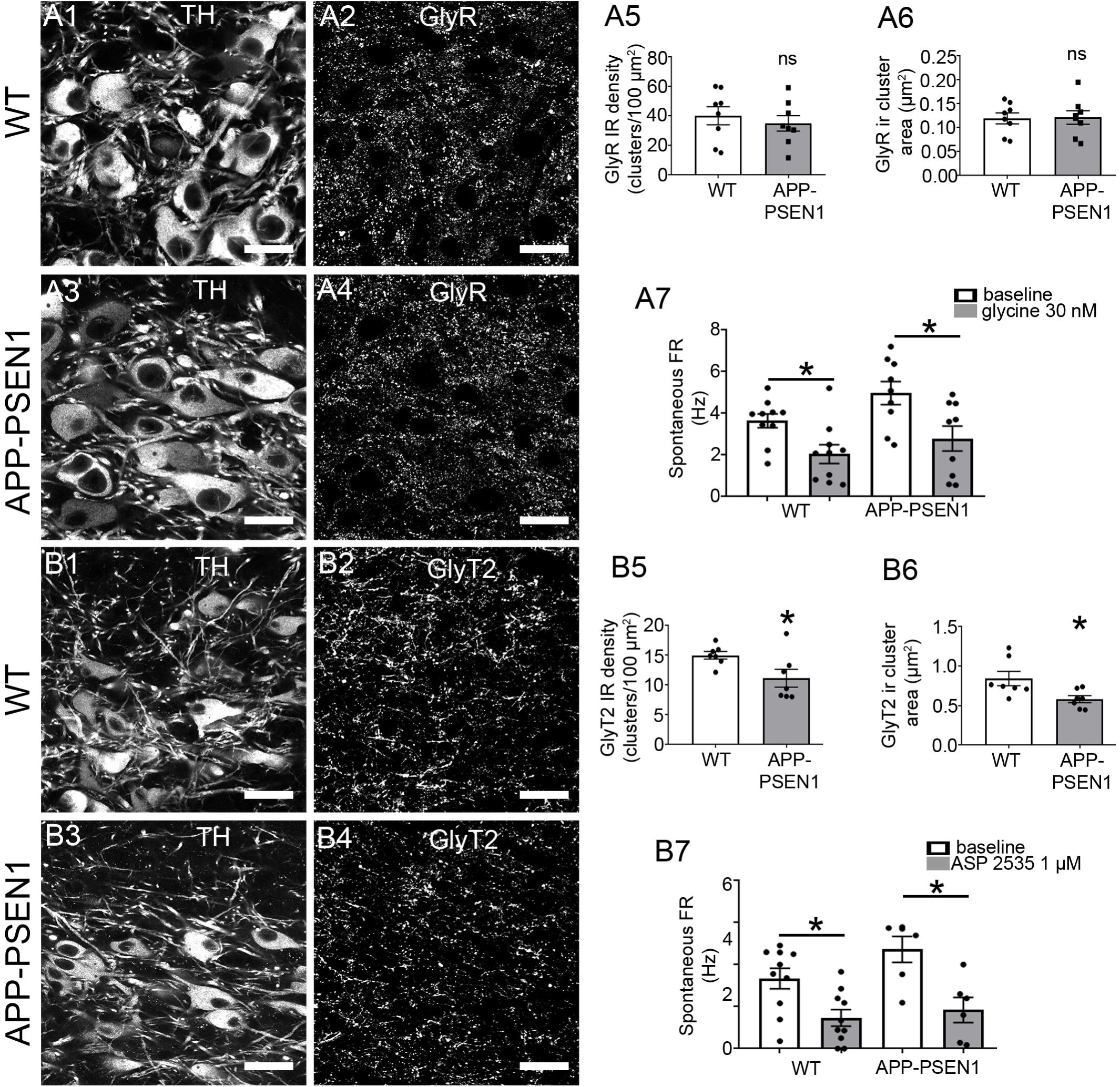
Quantification of the changes in the expression and function of GlyRs and GlyTs in LC neurons of the APP-PSEN1 mice. (A1) and (A2) show the pattern and intensity of TH and GlyR immunoreactivity, respectively, in tissue from a WT mouse. (A3) and (A4) show the comparative TH and GlyR immunoreactivity patterns, respectively, in in tissue from an APP-PSEN1 mouse. Note that samples for (A1-2) and (A3-4) were reacted and imaged under identical conditions. Quantification of GlyR immunoreactive cluster (A5) density and (A6) area. (A7) quantification of the comparative inhibitory effects of applied glycine (30 nM) on spontaneous LC FR in WT and APP-PSEN1 cells. (A5, 6) Two-tailed unpaired Student’s t test. (A7) Two way (genotype and drug) ANOVA. (B1) and (B2) show the pattern and intensity of TH and GlyT2 immunoreactivity, respectively, in tissue from a WT mouse. (B3) and (B4) show the comparative TH and GlyT2 immunoreactivity patterns, respectively, in in tissue from an APP-PSEN1 mouse. Note that samples for (B1-2) and (B3-4) were reacted and imaged under identical conditions. Quantification of GlyT2 immunoreactive cluster (B5) density and (B6) area. (B7) quantification of the comparative inhibitory effects of applied glycine transporter inhibitor ASP 1393 (1 µM) on spontaneous LC FR in WT and APP-PSEN1 cells. (B5, 6) Two-tailed unpaired Student’s t test. (B7) Two way (genotype and drug) ANOVA. The symbols represent the individual data points, the bars represent the means and the error bars the SEM. * = *P* < 0.05. Scale bars: 20 µm.

## Discussion

The study reveals for the first time, the expression profile of AβO in the LC in Alzheimer’s patients, one which is replicated in the APP-PSEN1 mouse model of increased Aβ production. Consequences of this elevated production of Aβ included an array of molecular changes involving immune, neurochemical and metabolic pathways, accompanied by neuronal hyperexcitability and altered NA brain levels. The data provide unique insights into the impact of AβO pathology in a brain region first to exhibit quintessential Alzheimer’s pathology.

### Importance of AβO in the LC in Alzheimer’s

The LC features prominently in assessments of the earliest pathological stages of Alzheimer’s, with the predominant characteristic being the presentation of tau tangles in this locus, prior to other brain regions (Weinshenker 2018). In stark contrast, the other post mortem pathological hallmark of Alzheimer’s, Aβ plaques, is largely overlooked as a contributor to the LC disease spectrum due to the relatively belated presentation of such pathology in this brain region, in most cases only at Braak stages V-VI (Braak et al. 2011). However, the importance of the Aβ system in contributing to the initial stages of Alzheimer’s could be a significant oversight due to recent evidence indicating that intraneuronal AβO accumulation precedes tau pathology in the entorhinal cortex, which alongside the LC, presents earliest with Alzheimer’s pathology (Welikovitch et al. 2018). Furthermore, compelling data indicate that AβO, rather than Aβ plaques impart the predominant toxic burden for this pathway (Ghag et al. 2018). More importantly, especially in terms of the disease phenotype in the LC, converging evidence points to a synergistic role between tau and AβO in mediating Alzheimer’s pathology (Crimins et al. 2013; Ittner and Gotz 2011; Spires-Jones and Hyman 2014). Indeed, AβO have been shown to enhance, whereas Aβ fibrils reduce, the aggregation of tau (Shin et al. 2019b). Furthermore, AβO promote the uptake of tau fibril seeds potentiating intracellular aggregation (Shin et al. 2019a) and mediate neurite degeneration (Jin et al. 2011). Mechanistic insights into this AβO-tau interaction arise from intriguing new data that demonstrate the involvement of NA, and α2_A_ adrenergic receptors (α2_A_ARs) in mediating tau hyperphosphorylation and Alzheimer’s-related pathology in human and animal samples (Zhang et al. 2020). Given the significant degree of LC axonal collaterals that provide negative feedback modulation of LC neuronal activity via α2_A_ARs (Aghajanian et al. 1977) (Cedarbaum and Aghajanian 1976), dysregulation of LC hyperexcitability, in combination with locally produced AβO, could set in motion the early levels of tau tangle formation distinctive in this brain region. Therefore, a systematic analysis of the relative temporal expression profiles for the presentation of tau tangle and AβO, rather than Aβ plaques, could provide a more insightful view into the early stages of Alzheimer’ disease.

Despite the recognised importance of the LC in Alzheimer’s, relatively little is known about the *functional* changes which occur in the constituent neurons of this nucleus. Whilst various brain imaging techniques have been used to examine the LC in patients at various stages of Alzheimer’s, most of these data relate to structural rather than functional changes (Betts et al. 2019; Peterson and Li 2018). The current study provides, to the best of our knowledge, the first analysis of the effects of Aβ-related pathology on the key functional determinant of LC-NA activity, namely spontaneous FR, demonstrating increased activity at the single cell level. This increased FR is likely to negatively impact on a range of behaviours, including cognitive performance (Usher et al. 1999), a heightened stress response (Abercrombie and Jacobs 1987) an increase in anxiety-like behaviour (McCall et al. 2015) and increased wakefulness (Carter et al. 2010). This in turn could help to elucidate the neurobiological basis for some of the key behavioural disturbances presented by early-stage Alzheimer’s patients, such as enhanced irritability, aggression and disturbances in sleep-wake cycles. Furthermore, if the increased LC FR evident in APP-PSEN1 mice proves to be a consistent feature in the early, pre-cognitive impairment stages of humans with the condition, this could support the further development of non-invasive early diagnostic strategies, such as those focused on the interaction between LC FR and pupillary dilation (Elman et al. 2017; Granholm et al. 2017).

The lack of an effect of Aβ-associated changes on the intrinsic membrane properties of LC neurons (Table 1), suggests that an alteration in the tone of neurotransmitter-receptor systems is the predominant basis for the enhanced tonic excitability. Whilst changes in the expression and function of neurotransmitter receptor systems may predominate in the early phases of Alzheimer’s-associated LC dysfunction, it is likely that this initial dysregulation in basal FR and consequent alteration in NA release, serves as the primer for the degenerative mechanisms which manifest in the decreased density of LC neurons at post mortem, and the deficits in cognitive (Jardanhazi-Kurutz et al. 2010) and immune (Heneka et al. 2010; Jardanhazi-Kurutz et al. 2011) functions mediated by NA. Importantly, some of these features have been demonstrated in this mouse model (Liu et al. 2008). Indeed, the level of LC activity regulates a key determinant of LC neuronal viability, namely oxidative stress, via ion channels which drive pacemaker activity (Sanchez-Padilla et al. 2014). In this study, evidence of oxidative stress was apparent in the LC, with a significant decrease detected in the APP-PSEN1 expression levels of SOD2, an enzyme involved in counteracting oxidative stress pathways. Given the significant evidence pointing to AβO mediating mitochondrial dysfunction (Leuner et al. 2012), including the attenuation of SOD2 levels (Cieslik et al. 2020), this observation suggests a considerable and consistent metabolic burden that needs to be maintained by these neurons, which may be the basis for their exaggerated decline in old age, leading to the impairments associated with cortical NA deficits in Alzheimer’s. Therefore, strategies to ameliorate this early stage neuronal hyperexcitability could have far reaching beneficial effects, not only on the whole LC-NA system, by limiting future cell loss and the disruptions to coordinated NA release, but also the main cortical sites for key dementia-related symptoms, because LC degeneration accelerates Alzheimer’s pathology in these brain regions (Heneka et al. 2010; Heneka et al. 2006; Jardanhazi-Kurutz et al. 2010; Jardanhazi-Kurutz et al. 2011; Rey et al. 2012).

### AβO and the LC inhibitory neurotransmitter systems

A striking finding was the significant impairment in the inhibitory effect of GABA on the tonic FR and the altered expression and function of α3-GABA_A_Rs of LC neurons in APP-PSEN1 mice. In light of the association of AβO-NU2 immunoreactivity with VGLUT2 signal, we assessed whether there were any changes in glutamatergic synaptic markers in APP-PSEN1 mice. There were no detectable changes in the levels of expression of presynaptic (VGLUT2) or postsynaptic (PSD-95) glutamatergic synaptic markers in APP-PSEN1 compared to WT mice (data not shown), suggesting a predominant impact of AβO on inhibitory synapse. Given this increased cytoplasmic and decreased membrane quotient for α3-GABA_A_Rs, the question therefore arises whether AβO-related pathology impairs its trafficking to the membrane, or enhances its internalisation from such domains. There is evidence for both mechanisms. The APP-PSEN1 model results in increased expression of amyloid precursor protein (APP) and production of Aβ (Borchelt et al. 1997), both of which could impact on the cytoplasmic trafficking and internalisation processes. Firstly, there is a strong interaction between AβO and GABA or GABA_A_Rs. The application of GABA down-regulates the endocytosis of AβO, thereby decreasing intracellular levels and AβO-induced cytotoxicity in WT mice (Sun et al. 2012). Furthermore, there is a bi-directional interaction between AβO and GABA_A_Rs. AβO impair GABA_A_R-mediated inhibition (Hu et al. 2017) by internalising GABA_A_Rs (Ulrich 2015). In contrast, chronic activation of GABA_A_Rs decreases Aβ_25-35_-mediated cytotoxicity (Lee et al. 2005).

An alternative explanation for the decreased α3 membrane expression could be an impairment in the trafficking of newly synthesised subunits to the membrane. GABA_A_Rs are assembled within the endoplasmic reticulum and transported to the Golgi apparatus for packaging onto vesicles (Comenencia-Ortiz et al. 2014). Subsequently, they are transported to the plasma membranes, from where they diffuse laterally into synapses (Jacob et al. 2005; Jacob et al. 2008; Thomas et al. 2005). Several proteins are involved in transporting the GABA_A_Rs from the Golgi, within transport vesicles, to the plasma membrane and their subsequent phosphorylation. One of these proteins, PLIC-1, facilitates the entry of GABA_A_R into the secretory pathway (Bedford et al. 2001). Importantly, PLIC-1 genetically associates with AD (Bertram et al. 2005) and regulates APP, as well as Aβ trafficking and processing (Hiltunen et al. 2006). Furthermore, the amount of PLIC-1 directly determines its influence on GABA_A_R trafficking since its overexpression promotes GABA_A_R membrane insertion (Bedford et al. 2001). Since the APP-PSEN1 mouse model is based on increased expression of APP, most probably beyond physiological levels, the expected increased demand for APP and Aβ handling by PLIC-1 in such mice could result in APP processing pathways predominating to the detriment of other systems, including GABA_A_Rs, resulting in a compromised delivery to plasma membranes. A robust counter-argument to this is that post-mortem samples from Alzheimer’s patients exhibited an almost indistinguishable LC α3-GABA_A_Rs expression profile to that observed in APP-PSEN1 samples. Indeed, in both mouse and human samples, a significant component of α3 immunoreactivity was located within cytoplasmic compartments, rather than the conventional enrichment on plasma membranes, suggesting that such altered trafficking of these receptors are a phenomenon of the disease itself, rather than the animal model; Finally, a more straightforward explanation could be AβO-mediated damage to the plasma membrane (Julien et al. 2018), which simply imposes space constraints for receptor insertion.

This strong association between Alzheimer’s-related pathology and α3-GABA_A_Rs is unsurprising. The α3 subunit links a number of brain regions which are known to be adversely affected especially in the early stages of the condition. These include monoaminergic brainstem centres (Corteen et al. 2015; Corteen et al. 2011; Gao et al. 1993) which degenerate with loss of their cortical modulatory presence, fear or stress circuitry such as the amygdala (Marowsky et al. 2012) and bed nucleus of the stria terminalis (Hortnagl et al. 2013), which most likely contribute to the neuropsychiatric symptoms, the olfactory system (Hortnagl et al. 2013) and the established impairment of olfaction (Daulatzai 2015), and the entorhinal cortex (Berggaard et al. 2018) and its role during the stage of mild cognitive impairment. A likely reason for the vulnerability of these α3-GABA_A_R centres could be the limited repertoire of GABA_A_R subtypes expressed by their constituent neurons. For example, compared to cortical pyramidal neurons, which express multiple α subunits and are able to compensate for the loss of individual subunits with limited consequences on neuronal excitability (Sur et al. 2001), any impairment in α3-GABA_A_R function is likely to have a tangible effect on the inhibitory modulation of these brain regions. Since many of these neurons already expend a heavy metabolic load in terms of their spontaneous activity and brain wide projections using unmyelinated axons, any impairment in their inhibitory regulation and consequent hyperexcitability is likely to hasten their demise, thereby accounting for their involvement in the AD processes at the earliest stages.

In light of the above, strategies to reverse any LC hyperexcitability due to impairments on GABA_A_Rs could not only address the associated symptoms, but also impede the progression of the condition, thereby enhancing the quality of life for those living with Alzheimer’s. Remarkably, GlyRs appeared impervious to Aβ-mediated pathology, even though, structurally, they belong to the same class of receptors as GABA_A_Rs. More importantly, we provide proof that the inhibition of GlyT1 reverses LC hyperexcitability in APP-PSEN1 mice. A number of compounds which inhibit the GlyT1 are in advanced stages of clinical testing as cognitive enhancers in schizophrenia (Cioffi 2018). The current study therefore provides the scientific rationale for the further exploration of such compounds, together with novel potent, and selective α3-GABA_A_R ligands, in the treatment of the different facets of cognitive and neuropsychiatric symptoms in Alzheimer’s patients.

## Acknowledgements

We gratefully acknowledge the contribution of the Alzheimer’s Society research monitors Chris Penrose, Jane Ward and Julie West in terms of their very constructive discussions throughout the duration of this project.

We would also like to thank the South West Dementia Brain Bank (SWDBB) for providing brain tissue for this study. The SWDBB is part of the Brains for Dementia Research programme, jointly funded by Alzheimer’s Research UK and Alzheimer’s Society (Brains for Dementia Research) and is supported by BRACE (Bristol Research into Alzheimer’s and Care of the Elderly) and the Medical Research Council.

The expert technical assistance of Stewart Gallacher, Daniel Arthur, Scott Rodaway, Angela Scutt, and Andrew Milner is gratefully acknowledged.

We are extremely grateful to Professor Arthur Butt (University of Portsmouth) for providing tissue from GFAP-GFP mice.

