## Supplementary material for "Identification of amyloid beta oligomers in locus coeruleus (LC) neurons of Alzheimer’s patients and their impact on LC oxidative stress, inhibitory neurotransmitter receptors and neuronal excitability": SI

**Supplementary Tables**

Table 1

Details of primary antibodies used

| Primary antibody | Host | Dilution | Source | Specificity/ Reference |
| --- | --- | --- | --- | --- |
| Amyloid beta. Recombinant monoclonal antibody corresponding to amino acids 1-42 of human beta amyloid (aa 4-10). This antibody recognises both human and murine Aβ. | Mouse | 1:3000 | Millipore MABN10 (clone WO-2) | (Ida et al. 1996) |
| Purified amyloid beta reactive to amino acid residues 17-24 of β amyloid | Mouse | 1:4000 | Biolegend (clone 4G8)  SIG-39220 | (Thal et al. 2002) |
| Amyloid beta oligomer specific-NU1 | Mouse | 1:5000 | William L. Klein  Northwestern University | (Lambert et al. 2007) |
| Amyloid beta oligomer specific-NU2 | Mouse | 1:5000 | William L. Klein  Northwestern University | (Lambert et al. 2007) |
| GABA_A_R α1  Antigen sequence α1N terminus amino acids 1–9  Rabbit # 21 ⁄ 7  Bleed # 04 ⁄ 10 ⁄ 1999 | Rabbit | 1:10,000 | A gift from Werner Sieghart, University of Vienna | Knockout mouse, (Corteen et al. 2011) |
| GABA_A_R α2  raised against amino acids 416-424 of the C-terminus | Rabbit | 1:500 | A gift from Werner Sieghart, University of Vienna | Knockout mouse, (Mitchell et al. 2018) |
| GABA_A_R α3  Synthetic peptide (aa 29 - 43 of rat GABA_A_R α3 subunit | Rabbit | 1:3000 | Synaptic Systems (224 303) | Knockout mouse, this study. |
| GABA_A_R β3  Fusion protein amino acids 370-433 of mouse GABA_A_R beta 3 subunit | Mouse | 1:2000 | Neuromab (75-149) clone (n87/ 25) | Knockout mouse, supplier |
| GABA_A_R γ2  Synthetic peptide (aa 39 - 67 of mouse GABA_A_R γ2 subunit | Rabbit | 1:4000 | Synaptic Systems (224003) | Labelling pattern as published with other antibodies |
| GABA_B_R | Mouse | 1:1000 | Neuromab clone N93A/49 | Knockout mouse, supplier |
| Gephyrin | Mouse | 1:1000 | Synaptic systems (147 021) | (Pfeiffer et al. 1984) and knockout mouse (Feng et al. 1998) |
| Glycine receptor  Synthetic peptide (aa 96-105) of GlyR α1 subunit. This antibody also recognises the β subunit | Mouse | 1:1000 | Synaptic Systems (146 011) | (Pfeiffer et al. 1984) |
| Glycine transporter 1 | Goat | 1:1000 | Santa Cruz (sc-16703) | (Jiang et al. 2007) |
| Glycine transporter 2 | Guinea pig | 1:500 | Frontier Institute (GlyT2-GP-Af800) | (Hondo et al. 2011) |
| IBA1 | Rabbit | 1:3000 | Wako (019-19741) | (Imai et al. 1996) |
| Superoxide 2/MnSOD | Rabbit | 1:2000 | Abcam (ab13534) | Labelling pattern as published with other antibodies |
| Tyrosine hydroxylase | Sheep | 1:3000 | Abcam, (ab113) | Raised to rat recombinant  TH. Labelling pattern as published with other antibodies |
| Tyrosine hydroxylase | Chicken | 1:2000 | Avēs labs (TYH) | Labelling pattern as published with other antibodies |
| VGAT | Goat | 1:3000 | Nittobo Medical  MSFR106130 | (Miura et al. 2006) |
| VGLUT2 | Guinea pig | 1:3000 | Nittobo Medical  MSFR106280 | Labelling pattern as published with other antibodies |

Table 2

Patient demographics for human tissue

| Diagnosis | BB NO | Brains for Dementia Research NO | MRC ID | AGE | SEX | Post mortem delay | HIST DIAGNOSIS | Braak tangle stage |
| --- | --- | --- | --- | --- | --- | --- | --- | --- |
| AD | 859 | B506 | BBN_4238 | 85 | F | 14 | AD definite, moderate arteriosclerotic small vessel disease, moderate to marked CAA | 5 |
| AD | 912 | B436 | BBN_14405 | 82 | F | 22 | AD definite, hippocampal sclerosis | 5 |
| AD | 920 | B409 | BBN_19615 | 75 | M | 7 | AD definite, moderate SVD, moderate CAA | 5 |
| AD | 946 | B346 |  | 93 | M | 30.5 | AD definite, moderately severe arteriosclerotic SVD | 5 |
| AD | 955 | B389 |  | 87 | F | 71.5 | AD definite, moderate SVD | 4 |
| Control | 881 | BC357 | BBN_4240 | 86 | F | 38.5 | Control, severe arteriosclerotic small vessel disease with microinfarcts | 1 |
| Control | 914 | BC536 | BBN_19608 | 96 | M | 21 | Control, no significant abnormalities | 2 |
| Control | 918 | BC542 | BBN_19613 | 85 | F | 13.5 | Control, microinfarct in hippocampus | 3 |
| Control | 927 | BC665 | BBN_19624 | 78 | M | 51.5 | Control, mild SVD | 2 |
| Control | 930 | BC736 | BBN_19627 | 94 | F | 29.5 | Control, moderate CAA | 2 |

**Supplementary Figures**

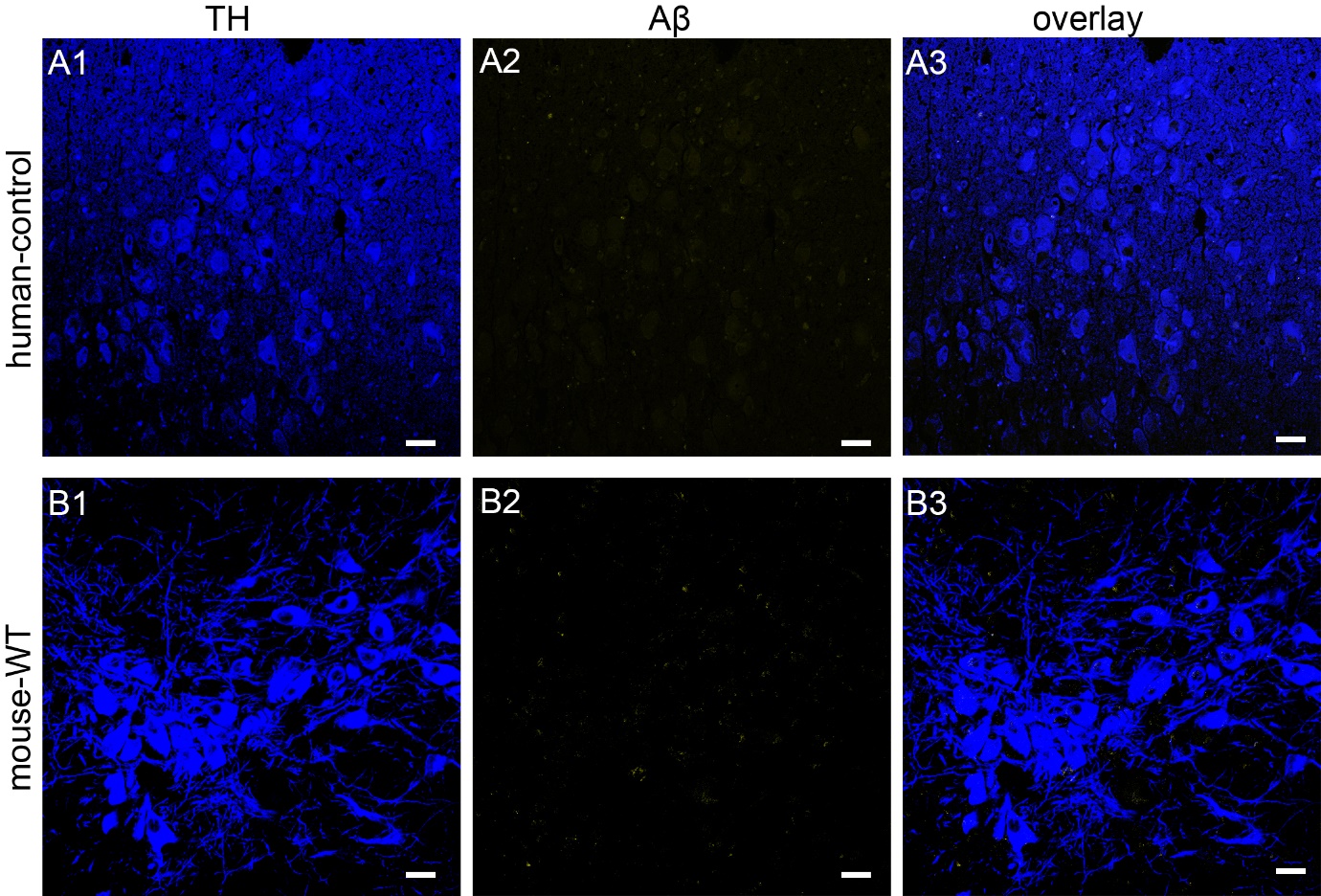

**Figure 1**

Confirmation of the specificity of Aβ immunoreactivity human control and mouse WT samples

(A) No specific Aβ signal was detected in the LC of human control samples.

(B) No specific Aβ signal was detected in the LC of WT mouse samples, processed and imaged under conditions identical to samples from APP-PSEN1 mice.

Scale bars: (A) 60 µm; (B) 30 µm

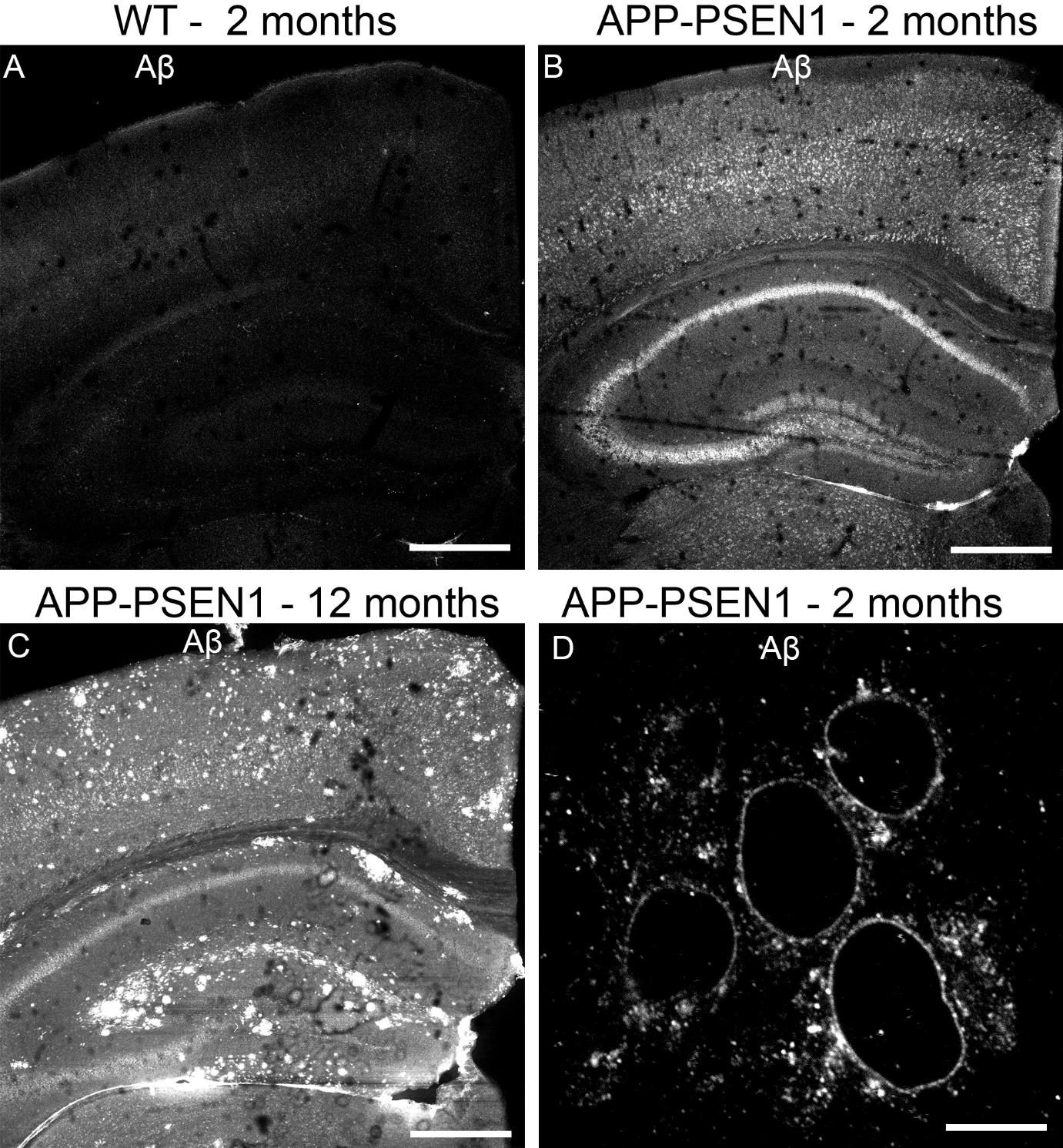

**Figure 2**

Demonstration of age-dependent amyloid β immunoreactivity pattern in cortical regions of APP-PSEN1 mice

(A) Immunoreactivity for Aβ was not detectable in tissue from WT mice aged 2 months or at any later stage (data not shown for old mice).

(B) shows Aβ immunoreactivity pattern at in tissue from a 2 month old APP-PSEN1 mouse, presenting as cytoplasmic signal throughout the hippocampus and neocortex. (C) in samples from 12 month old mice, immunoreactivity within neurons as well as extracellular plaques are evident.

(D) high resolution image of Aβ immunoreactivity in neocortical neurons highlighting the enrichment of signal in close location to plasma membranes, together with extracellular clusters.

Scale bars (A-C) 500 µm; (D) 10 µm.

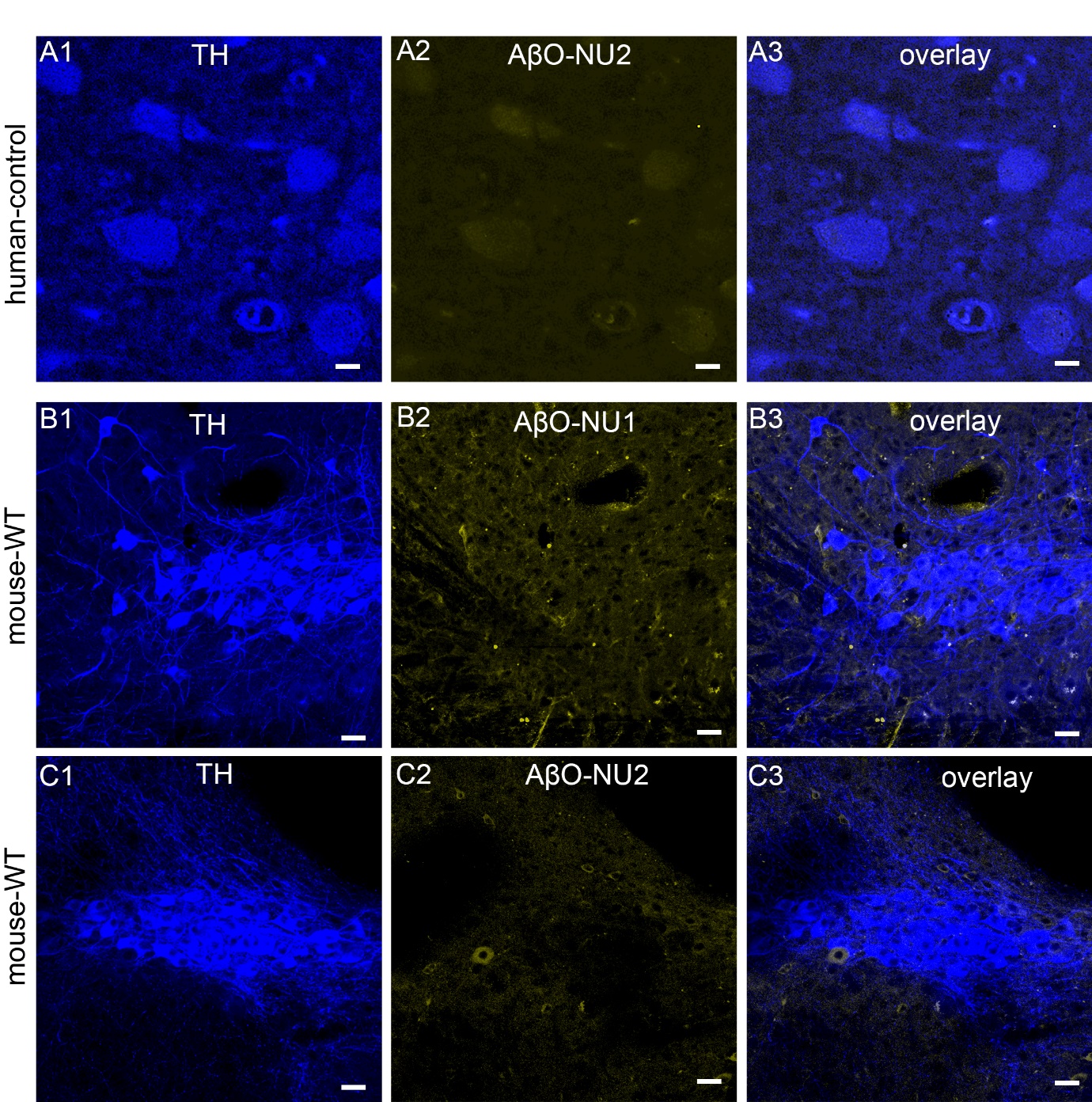

**Figure 3**

Confirmation of the specificity of AβO immunoreactivity in human control and mouse WT samples

(A) No specific AβO-NU2 signal was detected in the LC of human control samples.

(B) No specific AβO-NU1 signal was detected in the LC of WT mouse samples, processed and imaged under conditions identical to samples from APP-PSEN1.

(C) No specific AβO-NU2 signal was detected in the LC of WT mouse samples, processed and imaged under conditions identical to samples from APP-PSEN1.

Scale bars: (A) 20 µm; (B-C) 50 µm.

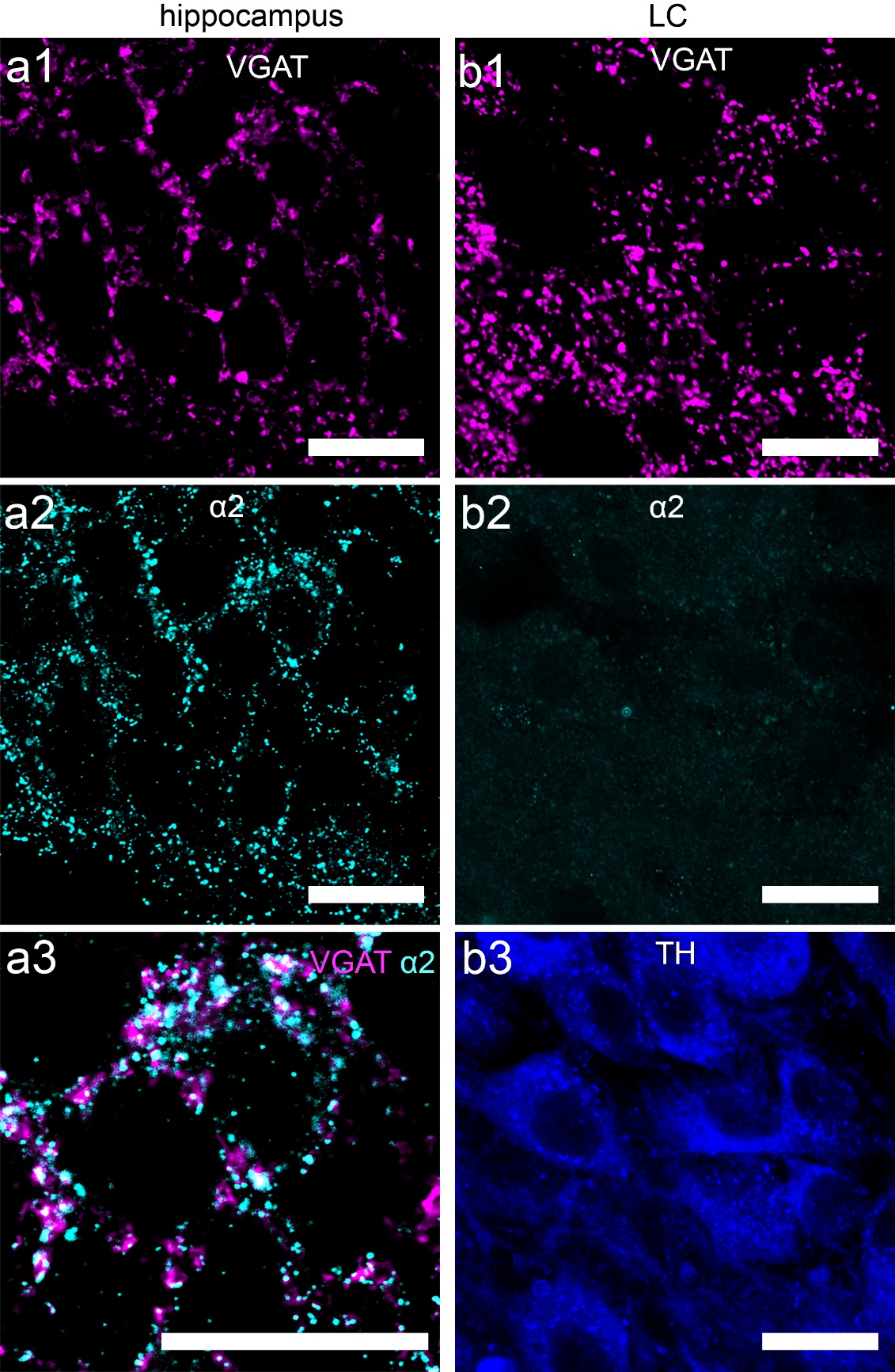

**Figure 4**

GABA_A_R α2 subunit is not detectable in the mouse locus coeruleus

(a) shows immunoreactivity for an antibody that recognises GABA_A_R α2 subunit in the mouse, in *stratum pyramidale* of CA1 hippocampus. The signal is closely associated with clusters immunopositive for VGAT, replicating the quintessential synaptic localisation pattern for this subunit. (b) using the same antibody, no specific signal was detectable in the LC.

Scale bars 20 µm.
